# Evolvability and constraint in the primate basicranium, shoulder, and hip and the importance of multi-trait evolution

**DOI:** 10.1101/2021.01.08.425985

**Authors:** Elizabeth R. Agosto, Benjamin M. Auerbach

## Abstract

The scapula shares developmental and functional relationships with traits of the basicranium, vertebral column, humerus, and clavicle. As a limb girdle, it also shares analogous characteristics with the pelvis. Despite these relationships, studies of primate shoulder evolution often focus on traits of the scapula in isolation. Such analyses may lead to spurious conclusions, as they implicitly model the scapula as evolving independent of other anatomical regions. Traits of the shoulder girdle share genetic covariances with each other, as well as potential covariances with dimensions of other skeletal elements. To create accurate models of shoulder evolution, it is imperative to account for the constraints imposed by these sources of covariance. Here, we use evolutionary quantitative methods to test a model in which shoulder morphological evolution is influenced by its developmental and functional covariances with the basicranium in the *Colobus* genus. This evolutionary relationship is also assessed with morphology of the pelvis to provide context to the evolutionary covariance among traits of the basicranium and shoulder girdle. Our results indicate potential evolutionary implications arising from covariances among the basicranium, shoulder, and pelvis. We further propose that the shoulder and basicranium may be examples of developmental, functional, and genetic covariances among traits that manifest an evolutionary suite of mutually constrained morphologies. We demonstrate novel evolutionary relationships among the shoulder girdle and basicranium that affect not only models of primate shoulder evolution but have broader implications for modeling trait evolution across the skeleton.

## Introduction

Traits within an organism often cannot evolve independent of other traits due to evolutionary covariance (Ackermann and Cheverud, 2000; Rolian et al. 2010; Grabowski and Roseman, 2015; Roseman and Auerbach 2015; Savell et al. 2016). This evolutionary covariance among traits arises from integration in response to genetic, developmental, or functional causes, among others (Felsenstein 1988; Cheverud 1996). The relationships among traits that ultimately result in their evolutionary covariance can be emergent through several mechanisms that all start at a genetic level (see Wagner 1996; Young 2004; Boughner 2016; Young et al. 2019). Despite the prevalence of evolutionary covariance, researchers often evaluate the evolution of traits in isolation. One example of this tendency is the study of primate shoulder evolution, where researchers often focus solely on traits of the scapula. The isolation of scapular measurements as a proxy for the overall functional and evolutionary morphology of the shoulder joint has been the precedent in primate studies for nearly a century (Schultz 1930; Schultz 1950; Ashton and Oxnard 1963; Oxnard 1963; Ashton and Oxnard 1964a; Ashton and Oxnard 1964b; Ashton et al. 1965; Oxnard 1967; Oxnard 1969; Roberts 1974; MacLatchy et al. 2000; Larson et al. 2007; Feuerriegel et al. 2017; Selby and Lovejoy 2017). However, the scapula shares functional relationships with many elements across the skeleton.

Traits exist within integrated groups, and so individual trait responses to selection are a combination of relationships with fitness independent of any other characteristic, and the correlated responses that trait has with other, genetically covarying traits (Rolian et al. 2010). These genetic correlations among traits can mask the strength and direction of natural selection. As a result, directional selection cannot be interpreted from the mean trait differences among populations (Lande and Arnold 1983; Houle 1991; Arnold 1992; Hansen and Houle 2008). This has been demonstrated in humans for the bones of the cranium (Katz et al. 2016), pelvis (Grabowski and Roseman 2015), hands and feet (Rolian et al. 2010), and body shape and size (Roseman and Auerbach 2015; Savell et al. 2016). As we note below, the scapula itself has developmental processes and shared functions that shape its morphology but are associated with other anatomical regions.

The scapula has functional relationships with the clavicle, humerus, basicranium, and vertebrae; the functional covariance between traits is the consequence of developmental processes (Cheverud 1996; Wagner and Altenberg 1996). Among the aforementioned skeletal elements, there are similar tissue contributions during ontogeny that shape these bones and their traits (Matsuoka et al. 2005; Huang et al. 2006; Durland et al. 2008; Valasek et al. 2010), such as post-otic neural crest (PONC), which migrates caudally from the cranium to form muscle attachments and regulate the mechanisms for muscle organization and connectivity between the head and shoulder (Matsuoka et al. 2005). Ultimately, these elements serve as attachment points for, and are therefore linked by, muscles responsible for movement at the shoulder, neck, and upper limb (Diogo and Wood 2012). Maintenance of shared developmental mechanisms would subsequently maintain the functional relationships among these anatomical regions. In addition to the shared development and function of these anatomical regions, the scapula is also analogous to the pelvis, as both girdles serve similar functions as the attachment point between the limbs and axial skeleton (Sears et al. 2015; Young et al. 2019). Sears and colleagues (2015) further demonstrate a lack of serial homology between the shoulder and pelvic girdles, as these structures do not share the same developmental processes. Rather, each displays its own unique developmental mechanisms and evolutionary history. Though the above cited research has explored shared developmental processes between the scapula and pelvis, such processes remain the subject of future studies. Therefore, how specific developmental processes and their underlying genomic causes relate to our questions are beyond the scope of this study. Nevertheless, genetic covariances underlie all of these relationships, and these covariances among traits potentially result in non-independent evolution of morphologies.

In the absence of experimental research design, developmental relationships are often difficult to ascertain from adult morphology, as these relationships become obscured through the superimposition of multiple determinants of covariance during ontogeny (Hallgrímsson et al. 2009; Castro et al. 2019; see also Huseynov et al. 2017). As such, adult morphology may be more reflective of the functional rather than the developmental relationships among traits. This can be problematic for ‘evo-devo’ based models of trait evolution. While we are unable to examine the developmental processes underlying evolutionary covariance among the scapula and other skeletal elements in this study, we know functional covariance among traits is emergent through developmental processes (Cheverud 1998). Thus, we focus on a functional argument which implicitly points to a developmental argument, but is not a developmental argument itself. Further, traits that are linked through shared function can impact organismal fitness by directly affecting the performance of a function, which may, therefore, have evolutionary consequences for these traits (Arnold 1983; Wainwright 1988), and thus the developmental processes which underlie them. The basicranium and the shoulder girdle are linked through mechanical function by the layers of connective and muscle tissue that physically connect these regions; these muscular relationships are fairly conserved among primates (Richmond et al. 2001; Diogo and Wood 2012).

Functional covariation is therefore important in shaping morphological evolution. The shoulder shares functions with adjacent anatomical regions, yet the evolutionary consequences of these relationships have not been explored in primates. To create accurate models of shoulder evolution, it is essential to account for the constraints imposed by the underlying covariances (i.e. developmental, genetic) between the shoulder and those skeletal elements in which it has functional relationships. To address this, we test a model in which the morphological evolution of the shoulder is influenced by its functional, and thus developmental, covariances with the basicranium. Specifically, we assess whether the ability of the shoulder girdle to respond to the force of directional selection would be constrained by its covariances with basicranial traits. We hypothesize that the developmental covariances underlying the functional relationships among the traits in these regions manifest an evolutionary and functional suite of mutually constrained morphologies.

To examine our model of non-independent shoulder evolution, we examine whether the functional and underlying developmental relationships between the scapula, clavicle, and basicranium are reflected in genetic covariances among their morphological traits that in turn, may constrain the ability of these traits to respond to selection. Two questions are asked: 1) Is the evolvability (Hansen and Houle 2008) of the shoulder with the basicranium smaller than estimated evolvabilities of the shoulder or basicranium with the pelvis? 2) Is the shoulder less evolvable when the basicranium or the pelvis is under stabilizing selection? To assess these questions, we obtained associated cranial and postcranial linear dimensions from a sample of 69 adults from the *Colobus* genus. Additional analyses on the same traits were performed on macaques (*Macaca mulatta*) and tamarins (*Saguinus oedipus* and *Saguinus fuscicollis illigeri*). We also include pelvic dimensions in our analysis, given the functional and developmental parallels between the shoulder and pelvic girdles, and to ascertain if there is a privileged potential evolutionary relationship between traits of the shoulder girdle and the basicranium compared with the shoulder and pelvis, or basicranium and pelvis (which are predicted to exhibit the most evolutionary independence based on both development and function). Measurements are mean-standardized prior to analysis. We calculated measures of evolvability and conditioned covariance following the equations of Hansen and Houle (Hansen and Houle 2008). Evolvability indices are estimated using 1000 simulated random selection gradients. Lower evolvability indicates greater evolutionary covariance among traits, and lower conditioned covariance indicates greater constraint among anatomical regions. Our results elucidate evolutionary relationships not previously explored and refine the model of shoulder evolution in primates, while suggesting broader considerations for the selection of traits to model trait evolution in all taxa.

## Materials and Methods

We measured all available adult *Colobus* genus specimens in North American museum collections with associated cranial and post-cranial skeletal elements necessary for this study, giving a total sample size of 69 individuals: *C. angolensis* (n=4), *C. guereza* (n=53), *C. polykomos* (n=9), and *C. satanas* (n=3). All individuals were adults, as indicated by fully fused epiphyses and spheno-occipital sutures; missing teeth in multiple individuals prevented the use of dental eruption for aging. Due to missing data, these samples are not sufficiently large enough by themselves to accurately estimate evolvability statistics. Pooling species was deemed reasonable as DNA analyses indicate the African colobine monkeys represent a monophyletic assemblage, with genus *Colobus* being more similar within itself than with the other two genera (*Piliocolobus and Procolobus*) in the Colobinae subfamily (Wang et al. 2012). Additionally, ANOVA results indicate no significant difference in the morphological traits among these species (p > 0.05; see Table S1). Lack of preservation of aspects of the crania and os coxa prevented the capture of certain measurements used in this study for several specimens. Due to the non-random nature of our missing data, missing values were not estimated (Allison 2002; Little and Rubin 2002). When the missing data were accounted for, our effective sample size on our pooled sample was reduced to a minimum of 37 and a maximum of 50 individuals. These sample sizes are adequate to accurately estimate evolvability statistics per the recommendation of Grabowski and Porto (2017), who specify n ≥ 36 with 10 traits, and n ≥ 37 with 15 traits.

Interlandmark distances for 21 dimensions from the basicranium, scapula, clavicle, and ilium (see Table S2) were calculated from three-dimensional coordinates (see Table S3 and Figure S1) captured using an Immersion G2x MicroScribe digitizer. Each coordinate was captured three, non-consecutive times by ERA and subjected to a repeated measures ANOVA (RMANOVA) to assess intra-observer error. Results of the RMANOVA indicated no significant difference among measurement rounds (p > 0.05), with an error of less than 1.2mm, so all dimensions were included. All interlandmark distances were mean-standardized within sex prior to all analyses.

Both males and females were included in all analyses. Differences due to sex were accounted for by applying a multivariate analysis of variance (MANOVA) and calculating the variance-covariance (VCV) matrix from the resulting residuals; henceforth, we refer to this as the sex-scaled VCV matrix.

Body size has been shown to affect how morphological traits respond to directional selection and can therefore affect estimates of evolvability and obscure non-size related evolutionary relationships among traits (Porto et al. 2013). To account for the effect of size on our analyses, we perform each analysis using the sex-scaled VCV matrix, and also a size-corrected VCV calculated using Equation 1 below. Following Porto and colleagues (2013), the size corrected VCV matrix (R) was obtained from the following relationship

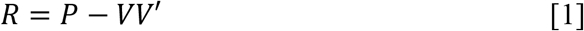

where P is the phenotypic VCV matrix, V is the size-related eigenvector, and V’ is its transpose.

Estimation of the VCV matrix can be difficult in biological systems due in part to issues with sample size or measurement error. This becomes increasingly difficult when matrix inversion is required (Twede and Heyden 2004; Marroig et al. 2012), and such inversions are required in the calculation of evolvability estimates. Small eigenvectors within the VCV matrix will reflect “noise” arising from measurement error more than meaningful variance, and thus transposes of these matrices will exaggerate the effects of the smallest eigenvectors. We control for noise in our VCV matrix estimates by locating the median of the variance of the second derivative derived from the absolute value of eigenvalues, removing all values below the median, replacing those values with the minimum eigenvalue, and recalculating the new VCV matrix using these new eigenvalues (Twede and Heyden 2004; Marroig et al. 2012). All analyses were performed on noise-controlled VCV matrices; we applied the correction to both the sex-scaled VCV matrix and R matrix calculated with Equation 1.

We used Hansen and Houle’s (2008) equations to calculate measures of evolvability and conditioned covariance among three anatomical regions: basicranium (skull), shoulder (scapula and clavicle), and pelvis (os coxa). Hansen and Houle’s methods are derived from Lande (1979) and colleagues’ (Lande and Arnold 1983) equation for the estimation of the average multivariate response to selection

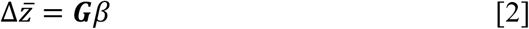

where 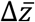 is the average response to selection, ***G*** is the genetic variance-covariance matrix, and *β* is the directional selection gradient. We used the within-group phenotypic variance-covariance (***P***) matrix as a substitute for the ***G*** matrix, as most studies indicate the ***G*** and ***P*** matrices are close to being proportional for morphological traits (Chverud 1988; Roff 1995; Roff 1996; Roseman 2012; Sodini et al. 2018).

Hansen and Houle’s (2008) approach uses the selection gradient to estimate multivariate evolvability. Within the context of Lande’s equation, evolvability is the capacity of a multivariate population mean to respond to the strength and direction of selection (Rolian 2009). Essentially, evolvability provides a measure of the potential for a group of traits to respond to directional selection; limitations to trait response to selection are imposed by the covariances among traits.

Greater evolutionary covariance among traits is indicated by lower measures of evolvability. Evolvability e(*β*) is calculated as

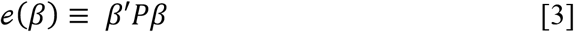

where variables are the same as in Equation 2, and *β*′ is the transpose of *β*.

The conditioned covariance provides a measure of the potential constraint a set of traits under stabilizing selection (x) imposes on a set of traits (y) to respond to directional selection. The conditioned covariance provides an estimate of the conditional genetic covariance of y given x (Hansen and Houle 2008) and can be thought of as a general measure of the genetic constraints among traits, or suites of traits. Conditioned covariance is calculated as

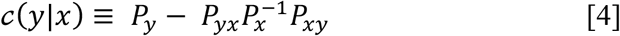

where *P*_*y*_ is the phenotypic variance in y, *P*_*xy*_ and *P*_*yx*_ are the column and row vectors of covariances between y and the traits in x, and *P*_*x*_ is the variance matrix of x (Hansen and Houle 2008; Hansen 2003; Hansen et al. 2003). In this equation, x denotes those traits being held constant by stabilizing selection, and y denotes traits under directional selection. Conditioned covariance is calculated for each pairing of anatomical regions, with each region acting as both x and y.

Five traits from each anatomical region (basicranium, shoulder girdle, and pelvic girdle) were used to minimize bias on our measures of evolvability. Thus, each analysis of evolvability used a total of fifteen interlandmark distances apportioned equally to the three anatomical regions. Indices of evolvability were calculated for the following groupings of anatomical regions: basicranium and shoulder girdle, shoulder and pelvic girdles, and basicranium and pelvic girdle. Analyzing groupings of anatomical regions allowed us to assess how suites of traits from different anatomical regions affect each other during evolution. Evolvability was also estimated among all three groupings combined, as our sample size is adequate to estimate this measure using 15 traits (n ≥ 37). To assess the effect of trait choice on our measures of evolvability, 13 different configurations of traits among regions were assessed. Traits that correlated highly with other traits *(r* > 0.8) (Table S4), were not included together in the same trait configurations for each analysis. Results on the following configuration of traits are presented in the main body of this paper: basicranium (NR, SCM, SNL, FMW, FML); shoulder (SSL, SBL, VB, GW, CLML); pelvis (AMIB, RAH, ASL, LIH, AW) (see Table 1 for measurement definitions and Figure 1 for their depiction on skeletal elements). All other trait configurations for each anatomical region and analysis can be found in the Supplementary Materials (see Table S5).

**Table 1.**
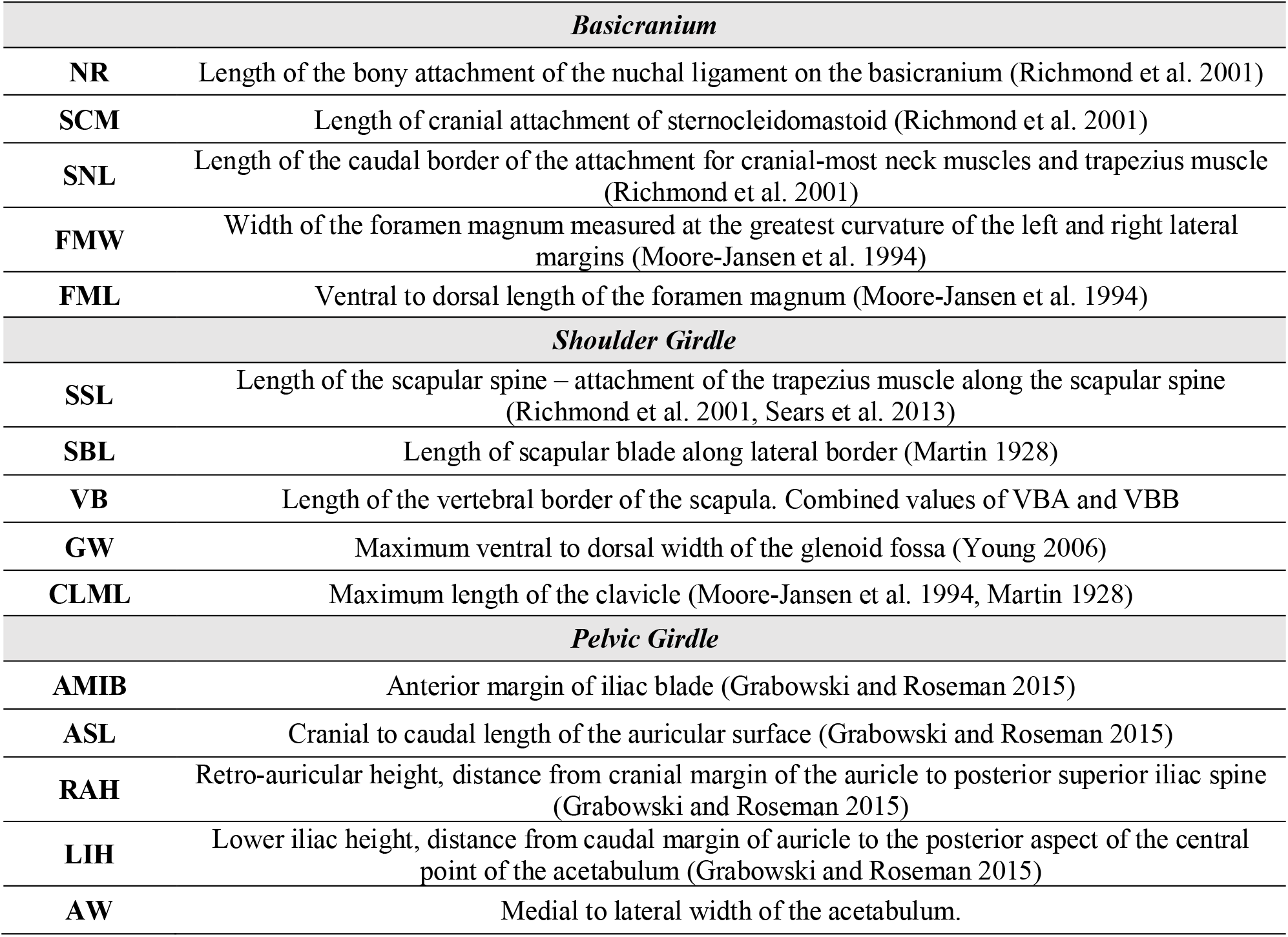
List of traits and trait descriptions for each anatomical region, along with cited sources of measurements.

| *Basicranium* |  |
| --- | --- |
| **NR** | Length of the bony attachment of the nuchal ligament on the basicranium (Richmond et al. 2001) |
| **SCM** | Length of cranial attachment of sternocleidomastoid (Richmond et al. 2001) |
| **SNL** | Length of the caudal border of the attachment for cranial-most neck muscles and trapezius muscle (Richmond et al. 2001) |
| **FMW** | Width of the foramen magnum measured at the greatest curvature of the left and right lateral margins (Moore-Jansen et al. 1994) |
| **FML** | Ventral to dorsal length of the foramen magnum (Moore-Jansen et al. 1994) |
| *Shoulder Girdle* |  |
| **SSL** | Length of the scapular spine – attachment of the trapezius muscle along the scapular spine (Richmond et al. 2001, Sears et al. 2013) |
| **SBL** | Length of scapular blade along lateral border (Martin 1928) |
| **VB** | Length of the vertebral border of the scapula. Combined values of VBA and VBB |
| **GW** | Maximum ventral to dorsal width of the glenoid fossa (Young 2006) |
| **CLML** | Maximum length of the clavicle (Moore-Jansen et al. 1994, Martin 1928) |
| *Pelvic Girdle* |  |
| **AMIB** | Anterior margin of iliac blade (Grabowski and Roseman 2015) |
| **ASL** | Cranial to caudal length of the auricular surface (Grabowski and Roseman 2015) |
| **RAH** | Retro-auricular height, distance from cranial margin of the auricle to posterior superior iliac spine (Grabowski and Roseman 2015) |
| **LIH** | Lower iliac height, distance from caudal margin of auricle to the posterior aspect of the central point of the acetabulum (Grabowski and Roseman 2015) |
| **AW** | Medial to lateral width of the acetabulum. |

**Figure 1.**
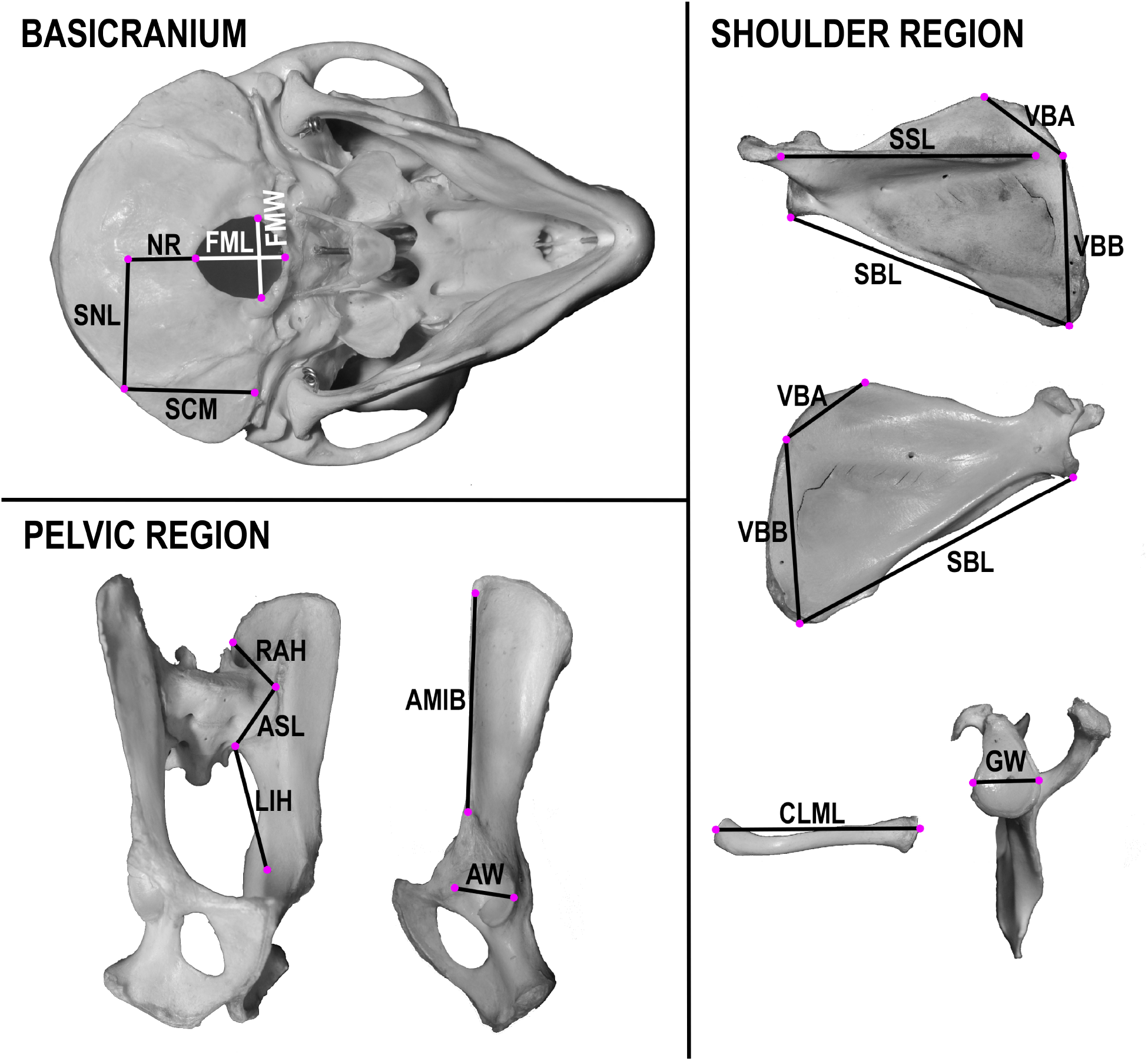
Linear dimensions collected from each anatomical region: caudal aspect of the basicranium; medial aspect of os coxa; lateral aspect of os coxa; dorsal aspect of scapula; ventral aspect of scapula; lateral aspect of scapula; cranial aspect of clavicle. All traits were measured from the left side. Refer to Table 1 for trait descriptions. Trait measurements are demonstrated on a macaque specimen and are homologous for colobines.

To calculate the average evolvability for each grouping of traits in phenotypic space, we subjected the phenotypic variance-covariance matrix to 1000 symmetrically distributed, randomly generated selection gradients. Selection gradients, which are vectors of partial regression coefficients of relative fitness on each trait in a multivariate system, summarize the direction and magnitude of selection acting on each trait (Lande and Arnold 1983; Hansen and Houle 2008; Rolian 2009). These selection gradients were generated using the ‘evolvability’ package by independently sampling each element of the selection gradient from a univariate, zero-mean Gaussian distribution with a unit variance and normalizing each vector by a unit length (Bolstad et al. 2014). 95% confidence intervals (CI’s) for both analyses were calculated by resampling each group of traits 1000 times with replacement, calculating the evolvability and conditioned covariance for each resampling, and deriving the 95% interval of the resulting distribution.

All analyses are performed in the R statistical environment (https://cran.r-project.org) using custom scripts and the ‘evolvability’ package (Bolstad et al. 2014). Both data and code are available upon request.

## Results

ANOVA results indicate no significant difference in trait dimensions (Table 2) between the four columbine species in the sample, however, there is a significant difference between the sexes within each species. Species were therefore pooled within each sex, and then measurements were mean-standardized within sex to allow all individuals to be analyzed together.

**Table 2.**
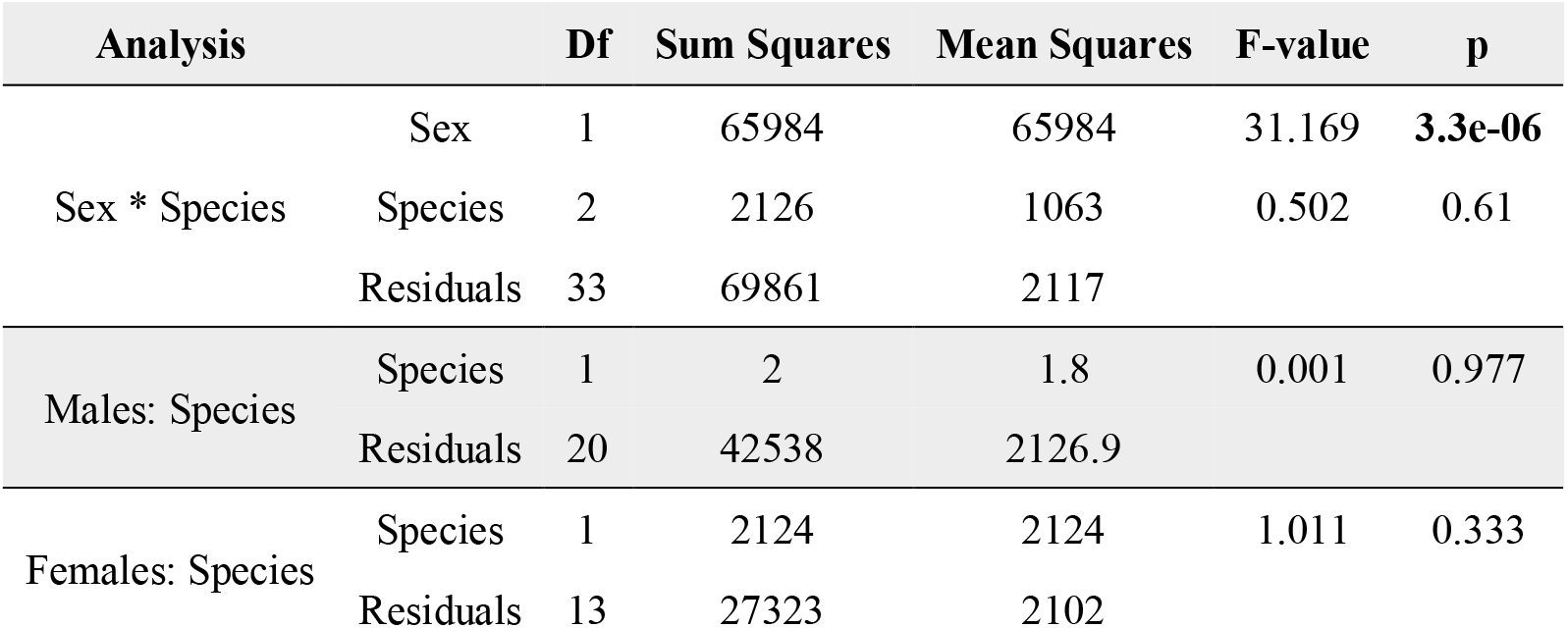
ANOVA results among the sexes and among species for each sex. Bold values are statistically significant (p < 0.05).

Traits were chosen to represent three anatomical regions: the basicranium, the shoulder girdle, and the pelvic girdle. These regions were placed into four groupings for the analysis of evolvability: all fifteen measures (basicranium, shoulder, and pelvis); measures of the shoulder girdle and the basicranium; measures of the shoulder girdle and the pelvic girdle; and measures of the basicranium and pelvic girdle. Table 3 presents the average estimated evolvability for each grouping of traits for both sex-scaled and size-corrected data. The mean estimated evolvability for all traits is between the highest and lowest evolvability estimates for pairwise trait groupings for both data treatments. This is expected, as this analysis incorporates the covariance among all anatomical regions. For both the sex-scaled and size-corrected data, the shoulder girdle – pelvic girdle grouping has the lowest estimate of evolvability. This indicates greater evolutionary covariance among these regions and less potential for them to independently respond to directional selection. For the sex-scaled data, the estimate of evolvability is lower for the basicranium – pelvic girdle grouping than the basicranium – shoulder girdle grouping. However, this relationship is reversed when the variance due to size is removed. The analysis on size corrected data shows that the basicranium – shoulder girdle is slightly less evolvable than the basicranium – pelvic girdle groupings. Our interpretation is that these regions have similar abilities to respond to directional selection. The 95% confidence intervals (CIs) among the three paired groupings of traits overlap, so while there is a pattern in how these three regions affect each other’s potential to respond to selection, it is not statistically significant.

**Table 3.**
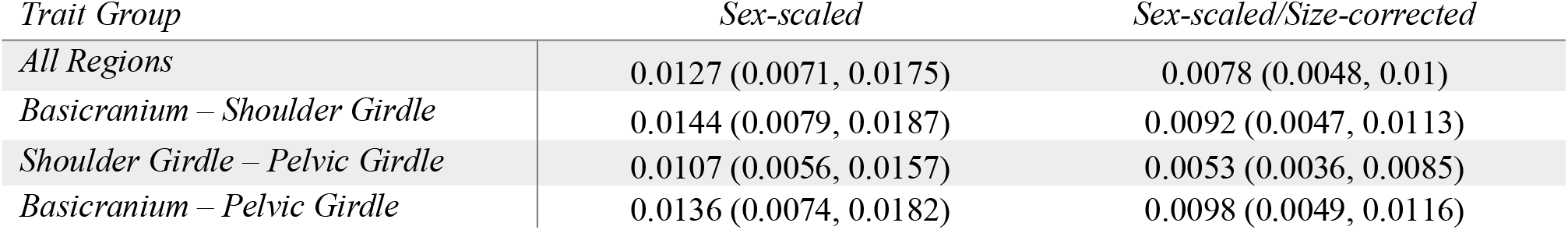
Mean evolvability and 95% confidence intervals among pairings of anatomical regions. Estimates for both sex-scaled and sex and size-scaled are presented.

| *Trait Group* | *Sex-scaled* | *Sex-scaled/Size-corrected* |
| --- | --- | --- |
| *All Regions* | 0.0127 (0.0071, 0.0175) | 0.0078 (0.0048, 0.01) |
| *Basicranium – Shoulder Girdle* | 0.0144 (0.0079, 0.0187) | 0.0092 (0.0047, 0.0113) |
| *Shoulder Girdle – Pelvic Girdle* | 0.0107 (0.0056, 0.0157) | 0.0053 (0.0036, 0.0085) |
| *Basicranium – Pelvic Girdle* | 0.0136 (0.0074, 0.0182) | 0.0098 (0.0049, 0.0116) |

Estimates of conditioned covariance were calculated for each pairwise grouping of anatomical regions used in the estimation of evolvability. We executed each pairwise analysis twice, with the traits in each anatomical region acting as if they were held constant by stabilizing selection (*x*) or those responding to directional selection (*y*) (see Materials and Methods for explanations of *x* and *y*), to assess whether one region imposes more constraint on another. The conditioned covariance among anatomical regions are presented in Table 4. The 95% CIs overlap for these analyses—meaning they are statistically the same—and so we focus on the patterns presented in the results. For both the sex-scaled and size-corrected data, we observe that the shoulder girdle is less evolvable if the pelvic girdle is under stabilizing selection than when the basicranium is under stabilizing selection. The pelvic girdle is less evolvable when the shoulder girdle, compared to the basicranium, is under stabilizing selection. The basicranium is less evolvable if the shoulder girdle is under stabilizing selection, compared to the pelvic girdle. We observe mutual constraint among anatomical regions, but the basicranium is more evolvable overall. Our results show that the basicranium is not constrained by the other two anatomical regions; rather, the basicranium imposes evolutionary constraint on both the shoulder and pelvic girdles. These results indicate the presence of hierarchical patterns of evolutionary constraint among anatomical regions.

**Table 4.** Mean conditioned covariance and 95% confidence intervals among anatomical regions. The anatomical region held constant by stabilizing selection is denoted by (x), whereas the anatomical region under directional selection is denoted by (y).

| *Trait Group* | *Sex-scaled* | *Sex-scaled/Size-corrected* |
| --- | --- | --- |
| Basicranium (x) – Shoulder Girdle (y) | 0.0523 (0.0186, 0.0772) | 0.0181 (0.0022, 0.0199) |
| Shoulder Girdle (x) – Basicranium (y) | 0.0760 (0.0254, 0.0998) | 0.0428 (0.0038, 0.0403) |
| Basicranium (x) – Pelvic Girdle (y) | 0.0462 (0.0172, 0.0742) | 0.0209 (0.0028, 0.0277) |
| Pelvic Girdle (x) – Basicranium (y) | 0.0780 (0.0315, 0.1027) | 0.0562 (0.0048, 0.051) |
| Shoulder Girdle (x) – Pelvic Girdle (y) | 0.0434 (0.0148, 0.0742) | 0.0184 (0.0028, 0.0255) |
| Pelvic Girdle (x) – Shoulder Girdle (y) | 0.0500 (0.0186, 0.0783) | 0.0135 (0.0025, 0.0252) |

As a subset of the total number of traits that were measured for each anatomical region (see Supplementary Material: Tables S2-3 and Figure S1) were used in these analyses, we repeated the estimation of evolvability and conditioned covariance using different compliments of traits within each anatomical region (Table S5). This was to test whether we introduced a bias in the traits we chose to include for each anatomical region. Such a bias would be revealed through inconsistencies in the results of these analyses. A similar trend in the patterns of evolvability and conditioned covariance is observed among all trait configurations (see Tables S6 and S7), suggesting these evolutionary relationships are not limited to only select traits from these anatomical regions. To further assess whether our results were isolated to the genus *Colobus*, we performed all analyses on two additional primate genera: *Macaca* and *Saguinus*. The results for these genera are consistent with, but not as pronounced, as those observed in *Colobus* (Tables S8 and S9). We also observe that the conditioned covariance among the basicranium and shoulder girdle is more similar in *Macaca*, suggesting mutual constraint among these regions.

## Discussion

Our results demonstrate potential evolutionary covariance among traits of the basicranium and shoulder girdle that result in their mutual evolutionary constraint. While the evolvability of the shoulder girdle is similarly constrained by the pelvic girdle, the demonstrated evolutionary covariance of the shoulder with the basicranium is unprecedented, even though generally overlooked developmental and functional relationships would anticipate such a relationship between these regions. Moreover, any evolutionary covariance between the cranium and postcranial elements in primates is heretofore unprecedented. The basicranium is not traditionally considered functionally or evolutionarily covariant with the shoulder girdle, as the base of the skull does not articulate with the scapula or clavicle. And yet, its evolution is tied to these other bones. Therefore, we provisionally answer our two questions raised in the Introduction: 1) the ability of the shoulder and basicranium together to respond to selection is at least as limited as the shoulder with the pelvis; and, 2) the shoulder is equally constrained to respond to selection by its underlying covariances with the basicranium and the pelvis, and vice versa. We have presented hypotheses throughout this paper that provide support to these findings, but the actual developmental mechanisms and processes that have led to these relationships are beyond the scope of this research. However, there is some evidence in the literature to set up testing developmental hypotheses for this relationship.

### Implications for the study of traits within evolutionary contexts

Our study’s implications for shared evolutionary constraint between the shoulder girdle and basicranium, especially in primates, has important applications to modeling the origins of variation among living taxa and for reconstructing evolutionary patterns from fossil remains. This analysis highlights the importance of considering the relation of traits on a genetic, developmental, and functional level simultaneously to model how they are likely to respond to evolution. Most crucial to the study of primate evolution, our results reemphasize that a level of caution should be taken when interpreting the evolution of individual traits in isolation, such as those in the shoulder girdle. For primates in general, including hominids, evolutionary change in one anatomical region will have related effects in areas that share developmental processes. For example, directional selection acting on the basicranium may have correlated effects on the scapula, though there was no selection acting directly on the scapula. The examination of the scapula or basicranium alone to understand the evolutionary forces that shaped either will fail to capture these evolutionary interactions. Moreover, within an anatomical region, as shown through other studies, individual traits within a functional complex will exhibit different directions and magnitudes of morphological change in response to the same selection, either through direct or correlated responses (Grabowski et al. 2010; Grabowski 2013; Savell et al. 2016). This phenomenon is visible in the fossil record as mosaic evolution (Schroeder et al. 2014), such as among traits that relate to hominid locomotion (Pilbeam 1996; Moyá-Solá et al. 2009; Nakatsukasa and Kunimatsu 2009; Feuerriegel et al. 2017). These relationships cannot be ascertained through the examination of changes in means of morphological traits—patterns of evolution—as these are not good indicators of the evolutionary processes that shaped them (Lande and Arnold 1983; Houle 1991; Arnold 1992; Hansen and Houle 2008; Grabowski et al. 2010; Grabowski 2013; Savell et al. 2016), or the underlying sources of covariation among traits (Hallgrímsson et al. 2009; Mitteroecker 2009). Detecting which traits were the target of selection, and which complexes of traits are more mutually constrained can only be assessed through a pan-primate comparison. The identification of *patterns* of morphological change within single anatomical regions, or single skeletal elements, and the interpretation of how those relate to functional differences among extinct primates are unaffected by our results and remain important for our understanding of primate evolution. However, our results call for extreme caution in ascribing evolutionary explanations to these patterns.

### Possible sources of mutual constraints in responses to selection among the shoulder, basicranium, and pelvis

The evolvability results for the limb girdles were expected. Due to the functional analogous nature of the limb girdles (e.g., Young et al. 2019), evolutionary relationships have previously been proposed for these regions (see Sears et al. 2015). These hypotheses tend to be driven by the similarity of general function of the limb girdles (i.e. anchoring the limbs to the torso), and the similarities in developmental genetic pathways (in addition to similar germ-cell layer origins). These relationships among the limb girdles may underlay the evolutionary relationship presented in the results of this study. It is also possible that some of the evolutionary covariance between these regions is also motivated by shared mechanical functions, as the species we chose to study are all quadrupeds.

However, as we note above, no such relationship has been anticipated previously between the basicranium and shoulder girdle. The estimated evolvabilities of the basicranium–shoulder girdle and shoulder girdle–pelvic girdle trait groupings are similar, and significantly lower than the evolvability of all traits combined. Thus, the shoulder girdle is mutually constrained by covariances with both the basicranium and the pelvic girdle, and vice versa, and more than the other two groupings we tested. However, we do not know whether the shoulder girdle is potentially more constrained by the basicranium or the pelvis, as our measures of evolvability and conditioned covariance are reporting slightly different patterns. While the question of whether the shoulder girdle is more mutually constrained by the basicranium or pelvic girdle remains unanswered, we do know that traits of the basicranium and shoulder girdle and traits of the limb girdles are equally labile in their ability to evolve.

We note that the sample size for these analyses is slightly smaller than recommended by Grabowski and Porto (2017) for measures of conditional evolvability, though we implemented methods to correct for possible biases in calculation of conditioned covariance. Without an adequate sample, measures of conditional evolvability (and thus likely conditioned covariance) estimates are negatively biased, which result in smaller estimated values and perceived stronger genetic constraint among traits (Grabowski and Porto 2017). It is therefore possible that our reported measures of conditioned covariance are biased downward, and therefore lower than the actual conditioned covariances among these traits. Nonetheless, we are confident in the conclusions we draw about mutual potential evolutionary constraint among the basicranium, shoulder and pelvis.

One additional confounding factor in our analysis is the effect of body size in shaping the results. Though we scaled by sex and individual measurement means prior to analyses, it is possible that some of the evolvability results we report were motivated by overall size. For this reason, we conducted an additional set of analyses to examine the evolvability of the three anatomical regions under consideration against five dimensions of the humerus (Table S10). While shared function links the humerus and the shoulder, the humerus shares no developmental or functional aspects with the basicranium and pelvis. We find that the highest evolvability estimates are associated with the humerus and basicranium, and the lowest are found between the humerus and shoulder. While this does not exclude the potential for size to affect our results, these additional results increase our confidence that the reported evolvability and conditioned covariance estimates reflect real evolutionary potentials.

### Future research directions: creating a testable framework for modeling evolutionary covariances with the shoulder

The outcomes of our study present a provocative potential for phenotypic and genomic relationships emergent within definitive anatomy that have not been predicted based on developmental models, but nevertheless may be reflected in functional relationships. The genomic and developmental mechanisms that link the traits we analyzed remain unknown; PONC are one of many hypothetical mechanisms that could underlie the covariance of the basicranium, scapula, and pelvis. Our findings, thus, await further verification within an experimental developmental model.

In the Introduction, we noted that emergent anatomical complexes arise from developmental, functional, and genetic evolutionary covariances. These could be referred to as modules (Wagner 1996; Schlosser and Wagner 2004; Wagner et al. 2007). However, we argue that modules are an imprecise concept within the framework of our study, as they are context-dependent and may be applied to multiple, nested levels of organization (cells, tissues, individuals, or populations, as summarized in Schlosser 2004). Finding a definition for modules and modularity that has consistent criteria by which to falsify their existence within an evo-devo framework remains elusive for the kind of anatomical complex we describe here. The unprecedented result showing potential equal mutual evolutionary constraint of the pelvis and basicranium on the shoulder may have further implications for creating a testable concept, which we refer to as a *functional trait complex*.

This concept is defined by traits wherein developmental, functional, and genetic covariances create an emergent anatomical complex in which traits mutually constrain independent responses to selection. Such complexes remain hypothetical but set up future studies that could seek shared regulatory genes or other genomic evidence for shared development, as well as studies incorporating the phylogenetic comparative method. Functional trait complexes would be limited and typified specifically by developmental and functional covariances that together yield evolutionary constraint. Developmental processes maintain functional associations among traits, and it is the integration of these two determinants of variation that affect how traits evolve (Hallgrímsson et al. 2009). We further propose this as the basis for testable criteria for one definition of modularity.

Models that predict how variation can be generated or structured through developmental processes can be used to make hypotheses of the patterns of morphological diversity and trait covariance (Young 2017). Previous studies have used this theoretical framework, in which traits that have shared developmental mechanisms and processes may exhibit evolutionary covariance, to assess serially homologous traits. This approach has been successfully applied to traits within the primate hands and feet (Rolian 2009; Rolian et al. 2010), limbs (as reviewed in Young 2017) and vertebrae (Williams and Russo 2015).

While the basicranium and shoulder girdle are not serial homologues, they do share a complex ontogenetic relationship involving both developmental mechanisms and mechanical function. These regions share several tissue contributions and genes during development, but most notably PONC forms attachment areas for muscles that span from the head to the shoulder girdle (Matsuoka et al. 2005). Developmental mechanisms shared among the basicranium and shoulder girdle would be involved in the functional relationship between these regions, and differences in the timing and/or duration of these developmental processes may affect the functional, and consequently morphological, variation observed among primate species. Among placental mammals, primates display the most variation in their shoulder morphology (Oxnard 1968; Preuschoft et al. 2010). The size, shape, and orientation of the muscles attaching to the scapula are a considerable source of variation in the primate shoulder girdle, as its morphological variation reflects the functional demands of the upper limb (Ashton et al. 1964; Oxnard 1968; Badoux 1974; Roberts 1974; Ashton et al. 1976; Young 2008; Preuschoft et al. 2010). The foundation of this variation is present at birth and is maintained through ontogeny (Young 2006). Expanding on this evidence, we hypothesize that the variance in shoulder morphology among primates may arise, in part, through the connectivity organized by PONC between the shoulder and adjacent anatomical regions to maintain functional relationships in the shoulder (Matsuoka et al. 2005). Due to myriad overlapping processes that contribute toward trait covariance, it is not possible to ascribe a single process as an explanation for the degree of morphological integration or the variance-covariance patterns among traits (Hallgrímsson et al. 2009), and development has been shown to be a robust, nonlinear process that may conceal the contributions of important genes and factors (Green et al. 2017). The resulting covariance structure among traits is most likely due to a combination of different causes that have yet to be described.

Our results suggest that this concept of a functional trait complex may be fruitfully assessed both for the shoulder and basicranium, as well as for the shoulder and pelvis. We realize that to successfully demonstrate the existence of a functional trait complex researchers would need to satisfactorily show that traits are linked developmentally and functionally, and that these have exerted evolutionary constraint. It is worth noting that approaches similar to the one we propose here have been published recently by Roseman and colleagues (2020), wherein the combination of development and morphological covariance enrich their mutual interpretation by combining evidence from both within the same study. The assessment of whether the concept of functional trait complexes is a productive avenue for evolution awaits future study, combining phylogenetic comparative methods with experimental developmental biology to test functional models of morphology.

## Conclusions

Our study adds novel evidence that sheds light on the complexities of shoulder evolution. We show that there is strong potential for misleading conclusions in models of shoulder evolution limited to traits of the scapula alone or ascribed to selection acting directly on specific trait morphologies. Rather, shoulder evolution is tied to the basicranium and the pelvis, as underlying covariances with both these regions constrain its evolution, and vice versa. The consistency of our results, regardless of the configuration of traits used, attests to the strength of the conclusions that can be drawn from our data. These results yield important insights for both functionally related traits and models of primate shoulder evolution, as well as implications for the interpretation of morphological changes in shoulder girdle variation within the fossil record. Caution should be exercised in any model of shoulder evolution that examines traits individually, as this implicitly assumes independent evolution. Morphological evolution is a whole-organism problem (Houle et al. 2010), and future models of shoulder evolution need to incorporate evolutionarily covariant traits.

## Declarations

### Funding

E.R.A was supported by National Science Foundation (NSF) Doctoral Dissertation Improvement Grant BCS – 1825995, the Thomas Fellowship through the University of Tennessee, and the College of Arts and Sciences at the University of Tennessee.

### Conflicts of Interest

Not applicable

### Availability of Data and Material

The dataset generated during and/or analyzed during the current study are available from the corresponding author on reasonable request.

### Code Availability

The custom code generated during the current study are available from the corresponding author on reasonable request.

## Acknowledgments

We thank C. Roseman, M. Grabowski, V. DeLeon, S. Williams, T. Capellini, and B. Hallgrímsson for helpful comments on this project. We would like to especially thank N. von Cramon-Taubadel and T. Weaver for pre-submission reviews on an early draft of this paper. E.R.A. thanks the many North American institutions for access to collections used in this study. E.R.A was supported by National Science Foundation (NSF) Doctoral Dissertation Improvement Grant BCS – 1825995, the Thomas Fellowship through the University of Tennessee, and the College of Arts and Sciences at the University of Tennessee.

## Notes

***Grant Sponsorship:*** This research is supported by a National Science Foundation Doctoral Dissertation Improvement Grant (NSF BCS-1825995).

### Competing Interest Statement

The authors have declared no competing interest.

## References

Ackermann, R.R., & Cheverud, J.M. (2000). Phenotypic covariance structure in tamarins (genus Saguinus): A comparison of variation patterns using matrix correlation and common principle component analysis. American Journal of Physical Anthropology 111, 489–501.

Ackermann, R.R., & Cheverud, J.M. (2002). Discerning evolutionary processes in patterns of tamarin (Genus Saguinus) craniofacial variation. American Journal of Physical Anthropology 117, 260–271.

Allison, P.D. (2002). Missing Data. Quantitative Applications in the Social Sciences, Volume 136. Newbury Park, CA: SAGE Publications.

Arnold, S.J. (1983). Morphology, performance, and fitness. American Zoologist 23, 347–361.

Arnold, S.J. (1992). Constraints on phenotypic evolution. American Naturalist 140, S85–S107.

Ashton, E.H., Flinn, R.M., Oxnard, C.E., & Spence, T.F. (1976). The adaptive and classificatory significance of certain quantitative features of the forelimb in primates. Journal of Zoology London 179, 515–556.

Ashton, E.H., & Oxnard, C.E. (1963). The musculature of the primate shoulder. Transactions of the Zoological Society of London 29, 554–650.

Ashton, E.H., & Oxnard, C.E. (1964). Oxnard, Functional adaptations in the primate shoulder girdle. Procedures of the Zoological Society of London 42, 49–66.

Ashton, E.H., & Oxnard, C.E. (1964). Locomotor patterns in primates. Procedures of the Zoological Society of London 142, 1–28.

Ashton, E.H., Oxnard, C.E., & Spence, T.F. (1965). Scapular shape and primate classification. Procedures of the Zoological Society of London 145, 125–142.

Badoux, D.M. (1974). An introduction to biomechanical principle in primate locomotion and structure. In F.A. Jenkins (Ed.), Primate Locomotion (pp. 1–44). New York: Academic Press.

Bolstad, G.H., Hansen, T.F., Pélabon, C., Falahati-Anbaran, M., Perez-Barrales, R., & Armbruster, W.S. (2014). Genetic constraints predict evolutionary divergence in Dalechampia blossoms. Philosophical Transactions of the Royal Society B. 369, 20130255.

Boughner, J.C. (2016). The tooth of the matter: the evo-devo of coordinated phenotypic change. In J. C. Boughner, C. Rolian (Eds.), Developmental Approaches to Human Evolution (pp. 35–60). New York: John Wiley & Sons.

Castro, J.P.L., Yancoskie, M.N., Machini, M., Belohlavy, S., Hiramatsu, L., Kučka, M., Beluch, W.H., Naumann, R. Skuplik, I., Cobb, J., Barton, N.H., Rolian, C., & Chan, Y.F. (2019). An integrative genomic analysis of the Longshanks selection experiment for longer limbs in mice. eLife 8, e42014. DOI: https://doi.org/10.7554/eLife.42014.

Cheverud, J.M. (1988). A comparison of genetic and phenotypic correlations. Evolution 42, 958–968.

Cheverud, J.M. (1996). Developmental integration and the evolution of pleiotropy. American Zoologist 36, 44–50.

Diogo, R., & Wood, B.A. (2012). Comparative Anatomy and Phylogeny of Primate Muscles and Human Evolution. Cleveland: C.R.C. Press.

Durland, J.L., Sferlazzo, M., Logan, M., & Burke, A.C. (2008). Visualizing the lateral somitic frontier in the Prx1Cre transgenic mouse. Journal of Anatomy 212, 590–602.

Felsenstein, J. (1988). Phylogenies and quantitative characters. Annual Review of Ecology and Systematics 19, 445–471.

Feuerriegel, E.M., Green, D.J., Walker, C.S., Schmid, P., Hawks, J., Berger, L.R., & Chuchill, S.E. (2017). The upper limb of Homo naledi. Journal of Human Evolution 104, 155–173.

Grabowski, M.W. (2013). Hominin obstetrics and the evolution of constraints. Evolutionary Biology 40, 57–75.

Grabowski, M. & Porto, A. (2017). How many more? Sample size determination in studies of morphological integration and evolvability. Methods in Ecology and Evolution 8, 592–603.

Grabowski, M., & Roseman, C.C. (2015). Complex and changing patterns of natural selection explain evolution of the human hip. Journal of Human Evolution 85, 94–110.

Green, R.M., Fish, J.L., Young, N.M., Smith, F.J., Roberts, B., Dolan, K., Choi, I., Leach, C.L., Gordon, P., Cheverud, J.M., Roseman, C.C., Williams, T.J., Marcucio, R.S., & Hallgrímsson, B. (2017). Developmental nonlinearity drives phenotypic robustness. Nature Communications 8, 1970.

Hallgrímsson, B., Jamniczky, H., Young, N.M., Rolian, C., Parsons, T.E., Boughner, J.C., & Marcucio, R.S. (2009). Deciphering the palimpsest: studying the relationship between morphological integration and phenotypic covariation. Evolutionary Biology 36, 355–376.

Hansen, T.F. (2003). Is modularity necessary for evolvability? Remarks on the relationship between pleiotropy and evolvability. Biosystems 69, 83–94.

Hansen, T.F., Armbruster, W.S., Carlson, M.L., & Pélabon, C. (2003). Evolvability and genetic constraint in Dalechampia blossoms: genetic correlations and conditional evolvability. Journal of Experimental Zoology B: Molecular Development and Evolution. 296B, 23–29.

Hansen, T.F., Houle, D. (2008). Measuring and comparing evolvability and constraints in multivariate characters. Journal of Evolutionary Biology 21, 1201–1219.

Houle, D. (1991). Genetic covariance of fitness correlates: What genetic correlations are made of and why it matters. Evolution 45, 630–648.

Houle, D., Govindaraju, D.R., & Hombolt, S. (2010). Phenomics: The next challenge. Nature Reviews Genetics 11, 855–866.

Huang, R., Christ, B., & Patel, K. (2006). Regulation of scapula development. Anatomy and. Embryology 211, S65–S71.

Huseynov, A., Ponce de León, M.S., & Zollikofer, C.P.E. (2017). Development of modular organization in the chimpanzee pelvis. Anatomical Record 300, 675–686.

Katz, D.C., Grote, M.N., Weaver, T.D. (2016). A mixed model for the relationship between climate and human cranial form. American Journal of Physical Anthropology 160, 593–603.

Lande, R. (1979). Quantitative genetic analysis of multivariate evolution, applied to brain: body size allometry. Evolution 33, 402–416.

Lande, R., & Arnold, S.J. (1983). The measurement of selection on correlated characters. Evolution 37, 1210–1226.

Larson, S.G., Jungers, W.L., Morwood, M.J., Sutikna, T., Jatmiko, Saptomo E.W., Due, R.A., & Djubiantono, T. (2007). Homo floresiensis and the evolution of the hominin shoulder. Journal of Human Evolution 53, 718–731.

Little, R.J.A., & Rubin, D.B. (2002). Statistical Analysis with Missing Data. Second Edition New York: John Wiley & Sons.

MacLatchy, L., Gebo, D., Kityo, R., & Pilbeam, D. (2000). Postcranial functional morphology of Morotopithecus bishopii, with implications for the evolution of modern ape locomotion. Journal of Human Evolution 39, 159–183.

Marroig, G., Melo, D.A.R., & Garcia, G. (2012). Modularity, noise, and natural selection. Evolution 66, 1506–1524.

Martin, R. (1928). Lehrbuch der Anthropologie in Systematischer Darstellung mit Besonderer Berücksichtigung der Anthropologischen Methoden für Studierende Ärtze und Forschungsreisende. Zweiter Band: Kraniologie, Osteologie. Second Edition. Jena: Gustav Fischer.

Matsuoka, T., Ahlberg, P.E., Kessaris, N., Iannarelli, P., Dennehy, U., Richardson, W.D., McMahon, A.P., & Koentges, G. (2005). Neural crest origins of the neck and shoulder. Nature 436, 347–355.

Mitteroecker, P. (2009). The developmental basis of variational modularity: insights from quantitative genetics, morphometrics, and developmental biology. Evolutionary Biology 36, 377–385.

Moore-Jansen, P.M., Ousley, S.D., & Jantz, R.J. (1994). Data Collection Procedures for Forensic Skeletal Material. Report of Investigations No. 48. Knoxville, TN: The University of Tennessee.

Moyá-Solá, S., Köhler, M., Alba, D.M., Casanovas-Vilar, I., & Galindo, J. (2009). Peirolapithecus catalaunicus, a new Middle Miocene great ape from Spain. Science 306, 1339–1344.

Nakatsukasa, M., & Kunimatsu, Y. (2009). Nacholapithecus and its importance for understanding hominoid evolution. Evolutionary Anthropology. 18, 103–119 (2009).

Oxnard, C.E. (1963). Locomotor adaptations in the primate forelimb. Symposium of the Zoological Society of London 10, 165–182.

Oxnard, C.E. (1967). The functional morphology of the primate shoulder as revealed by comparative anatomical, osteometric, and discriminant function techniques. American Journal of Physical Anthropology 26, 219–240.

Oxnard, C.E. (1968). The architecture of the shoulder in some mammals. Journal of Morphology 126, 249–290.

Oxnard, C.E. (1969). Evolution of the human shoulder: some possible pathways. American Journal of Physical Anthropology 30, 319–332.

Pilbeam, D. (1996). Genetic and morphological records of the Hominoidea and hominid origins: A synthesis. Molecular Phylogenetics and Evolution 5, 155–168.

Pilbeam, D., & Young, N.M. (2004). Hominoid evolution: Synthesizing disparate data. Comptes Rendus Palevol 3, 305–321.

Porto, A., Shirai, L.T., de Oliveira, F.B., & Marroig, G. (2013). Size variation, growth strategies and the evolution of modularity in the mammalian skull. Evolution 67, 3305–3322.

Preuschoft, H., Hohn, B., Scherf, H., Schimdt, M., Krause, C., & Witzel, U. (2010). Functional analysis of the primate shoulder. International Journal of Primatology 31, 301–320.

Richmond, F.J.R., Singh, K., & Corneil, B.D. (2001). Neck Muscles in the Rhesus Monkey. I. Muscle Morphometry and Histochemistry. Journal of Neurophysiology 86, 1717–1728.

Roberts, D. (1974). Structure and function of the primate scapula. In F.A. Jenkins (Ed.), Primate Locomotion (pp. 171–200). New York: Academic Press.

Roff, D.A. (1995). The estimation of genetic correlations from phenotypic correlations: a test of Cheverud’s conjecture. Heredity 74, 481–490.

Roff, D.A. (1996). The evolution of genetic correlations: an analysis of patterns. Evolution 50, 1392–1403.

Rolian, C. (2009). Integration and evolvability in primate hands and feet. Evolutionary Biology 36, 100–117.

Rolian, C., Lieberman, D.E., & Hallgrímsson, B. (2010). The coevolution of human hands and feet. Evolution 64, 558–1568.

Roseman, C.C. (2012). The soundness of the Cheverud Conjecture and its implications for the study of human evolution. American Journal of Physical Anthropology 147, 252.

Roseman, C.C., & Auerbach, B.M. (2015). Ecogeography, genetics, and the evolution of human body form. Journal of Human Evolution 78, 80–90.

Roseman, C.C., Capellini, T.D., Jagoda, E., Williams, S.A., Grabowski, M., O’Connor, C., Polk, J.D., Cheverud, J.M. (2020). Variation in mouse pelvic morphology maps to locations enriched in Sox9 Class II and Pitx1 regulatory features. Journal of Experimental Zoology B: Molecular and Developmental Evolution 334:100–112.

Savell, K.R.R., Auerbach, B.M., & Roseman, C.C. (2016). Constraint, natural selection, and the evolution of human body form. Proceedings of the National Academy of Sciences USA 113, 9492–9497.

Schlosser, G. (2004). The role of modules in development and evolution. In G. Schlosser & G.P. Wagner (Eds.), Modularity in Development and Evolution (pp. 519-582). Chicago: University of Chicago Press.

Schlosser, G., & Wagner, G.P. (2004) Introduction: the modularity concept in developmental and evolutionary biology. In G. Schlosser & G.P. Wagner (Eds.), Modularity in Development and Evolution (pp. 1-16). Chicago: University of Chicago Press.

Schultz, A.H. (1930). The skeleton of the trunk and limbs of higher primates. Human Biology 2, 303–438.

Schultz, A.H. (1950). The physical distinctions of man. Proceedings of the American Philosophical Society 94, 428–449.

Schroeder, L., Roseman, C.C., Cheverud, J.M., & Ackermann, R.R. (2014). Characterizing the evolutionary path(s) to early Homo. PLoS ONE 9, e114307.

Sears, K.E., Bianchi, C., Powers, L., & Beck, A.L. (2013). Integration of the mammalian shoulder girdle within populations and over evolutionary time. Journal of Evolutionary Biology 26, 1536–1548.

Sears, K.E., Capellini, T.D., & Diogo, R. (2015). On the serial homology of the pectoral and pelvic girdles of tetrapods. Evolution 69, 2543–2555.

Selby, M.S., & Lovejoy, C.O. (2017). Evolution of the hominoid scapula and its implications for earliest hominid locomotion. American Journal of Physical Anthropology 162, 682–700.

Sodini, S.M., Kemper, K.E., Wray, N.R., & Trzaskowski, M. (2018). Comparison of genotypic and phenotypic correlations: Cheverud’s conjecture in humans. Genetics 209, 941–948.

Twede, D.R., & Hayden, A.F. (2004). Refinement and generalization of the extension method of covariance matrix inversion by regularization for spectral filtering optimization. Proc. SPIE 5159, Imaging Spectrometry IX. doi: 10.1117/12.506993.

Valasek, P., Theis, S., Krejci, E., Grim, M., Maina, F., Shwartz, Y., Otto, A., Huang, R., & Patel, K. (2010). Somatic origin of the medial border of the mammalian scapula and its homology to the avian scapula blade. Journal of Anatomy 216, 482–488.

Wagner, G.P. (1996). Homology, natural kinds, and the evolution of modularity. American Zoologist 36, 36–43.

Wagner, G.P., & Altenberg, L. (1996) Complex adaptations and the evolution of evolvability. Evolution 50, 967–976.

Wagner, G.P., Pavlicev, M., & Cheverud, J.M. (2007). The road to modularity. Nature Reviews Genetics 8, 921–931.

Wainwright, P.C. (1988). Morphology and ecology: the functional basis of feeding constraints in Caribbean labrid fishes. Ecology 69, 635–645.

Wang, X.P., Yu, L., Roos, C., Ting, N., Chen, C.P., Wang, J., & Zhang, Y.P. (2012). Phylogenetic relationships among the colobine monkeys revisited: new insights from analyses of complete mt genomes and 44 nuclear non-coding markers. PLoS ONE 7, e36274.

Williams, S.A. & Russo, G.A. (2015). Evolution of the Hominoid vertebral column: the long and short of it. Evolutionary Anthropology 24, 15–32.

Young, M., Selleri L., & Capellini, T.D. (2019). Genetics of scapula and pelvis development: an evolutionary perspective. Current Topics in Developmental Biology 132, 311–349.

Young, N.M. (2003). A reassessment of living hominoid postcranial variability: implications for ape evolution. Journal of Human Evolution 45, 441–464.

Young, N.M. (2004). Modularity and integration in the Hominoid scapula. Journal of Experimental Zoology Part B: Molecular and Developmental Evolution 302B, 226–240 (2004).

Young, N.M. (2006). Function, ontogeny and canalization of shape variance in the primate scapula. Journal of Anatomy 209, 623–636.

Young, N.M. (2008). A comparison of the ontogeny of shape variation in the Anthropoid scapula: functional and phylogenetic signal. American Journal of Physical Anthropology 136, 247–264.

Young, N.M. (2017). Integrating “Evo” and “Devo”: The limb as a model structure. Integrative and Comparative Biology 57, 1293–1302.

