## Supplementary material for "Evolvability and constraint in the primate basicranium, shoulder, and hip and the importance of multi-trait evolution": SI_Appendix

**This PDF file includes:**

Fig. S1
Tables S1 to S10
References for SI reference citations

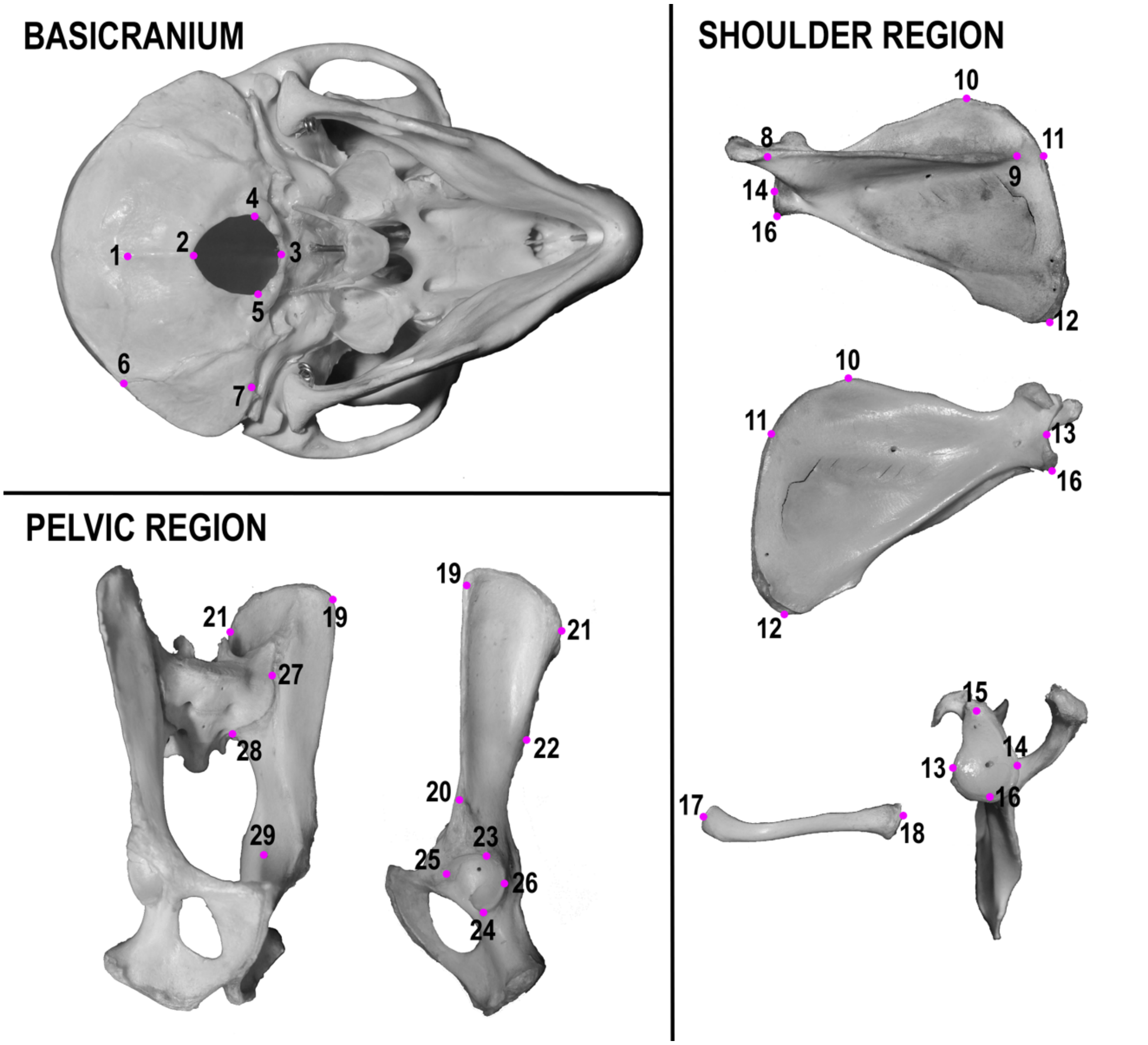


**Figure S1.** Landmarks taken from each anatomical region analyzed: caudal aspect of the basicranium; medial aspect of articulated pelvis; lateral aspect of left os coxa; dorsal aspect of scapula; ventral aspect of scapula; lateral aspect of scapula; cranial aspect of clavicle. All landmarks were taken from the left side. Refer to Table S3 for list of landmarks and descriptions. Landmark locations are demonstrated on a macaque specimen and are homologous for colobines.

**Table S1.** ANOVA results among the sexes and among species for each sex. Bold values are statistically significant ($p < 0.05$).

| Analysis | | Df | Sum Squares | Mean Squares | F-value | p |
|---|---|---|---|---|---|---|
| Sex * Species | Sex | 1 | 65984 | 65984 | 31.169 | **3.3e-06** |
| | Species | 2 | 2126 | 1063 | 0.502 | 0.61 |
| | Residuals | 33 | 69861 | 2117 | | |
| Males: Species | Species | 1 | 2 | 1.8 | 0.001 | 0.977 |
| | Residuals | 20 | 42538 | 2126.9 | | |
| Females: Species | Species | 1 | 2124 | 2124 | 1.011 | 0.333 |
| | Residuals | 13 | 27323 | 2102 | | |

**Table S2.** Linear measures calculated from landmarks, measurement descriptions, and descriptive statistics for each measurement for both raw (in mm) and scaled measures. The mean and standard deviation is shown by sex for raw measures and with sexes combined for scaled measures.

| Trait | Landmarks | Description | Females | | Males | | Scaled | |
|---|---|---|---|---|---|---|---|---|
| | | | Mean | SD | Mean | SD | Mean | SD |
| ***Basicranium*** | | | | | | | | |
| NR | 1 – 2 | Length of the bony attachment of the nuchal ligament on the basicranium (1). | 17.43 | 2.51 | 20.26 | 2.81 | 1.00 | 0.15 |
| SCM | 6 – 7 | Length of cranial attachment of sternocleidomastoid (1). | 18.33 | 1.58 | 21.76 | 2.65 | 1.00 | 0.11 |
| SNL | 1 – 6 | Length of the caudal border of the attachment for cranial-most neck muscles and trapezius muscle (1). | 15.39 | 3.05 | 17.99 | 2.84 | 1.00 | 0.19 |
| FMW | 4 – 5 | Width of the foramen magnum measured at the greatest curvature of the left and right lateral margins (2). | 15.16 | 0.77 | 16.03 | 1.00 | 1.00 | 0.06 |
| FML | 2 – 3 | Ventral to dorsal length of the foramen magnum (2). | 15.72 | 1.21 | 15.73 | 1.00 | 1.00 | 0.07 |
| ***Shoulder Girdle*** | | | | | | | | |
| SSL | 8 – 9 | Length of the scapular spine – attachment of the trapezius muscle along the scapular spine (1,3). | 63.43 | 5.77 | 73.05 | 6.98 | 1.00 | 0.09 |
| SBL | 12 – 16 | Length of scapular blade along lateral border (4). | 72.68 | 5.94 | 83.73 | 5.96 | 1.00 | 0.09 |
| SBW | 11 – 14 | Maximum medial to lateral breadth of scapular blade (2,4). | 63.23 | 4.91 | 72.22 | 4.53 | 1.00 | 0.07 |
| SBH | 10 – 12 | Cranial to caudal height of scapular blade (2,4). | 68.62 | 6.52 | 80.1 | 7.39 | 1.00 | 0.10 |
| VB | 10 – 11<br>11 – 12 | Length of the vertebral border of the scapula. Combined values of VBA (10-11) and VBB (11-12). | 78.11 | 8.29 | 91.55 | 9.57 | 1.00 | 0.11 |
| GW | 13 – 14 | Maximum ventral to dorsal width of glenoid fossa (5). | 10.73 | 0.86 | 11.66 | 1.13 | 1.00 | 0.09 |
| GL | 15 – 16 | Maximum cranial to caudal length of the glenoid fossa (5). | 15.65 | 1.22 | 17.01 | 1.34 | 1.00 | 0.08 |
| CLML | 17 – 18 | Maximum length of the clavicle (2,4). | 53.32 | 4.81 | 63.01 | 4.66 | 1.00 | 0.09 |
| ***Pelvic Girdle*** | | | | | | | | |
| AMIB | 19 – 20 | Anterior margin of iliac blade (6). | 59.54 | 7.89 | 60.67 | 9.26 | 1.00 | 0.14 |
| PMIB | 21 – 22 | Posterior margin of iliac blade (6). | 42.81 | 4.37 | 49.04 | 3.91 | 1.00 | 0.09 |
| ASL | 27 – 28 | Cranial to caudal length of the auricular surface (6). | 23.24 | 3.6 | 25.66 | 4.23 | 1.00 | 0.16 |
| RAH | 21 – 27 | Retro-auricular height, distance from cranial margin of the auricle to posterior superior iliac spine (6). | 37.17 | 4.27 | 42.11 | 3.19 | 1.00 | 0.10 |
| LIB | 19 – 27 | Lateral iliac breadth, distance from cranial margin of the auricle to the anterior superior iliac spine (6). | 50.7 | 6.46 | 56.31 | 3.59 | 1.00 | 0.11 |
| LIH | 28 – 29 | Lower iliac height, distance from caudal margin of auricle to the posterior aspect of the central point of the acetabulum (6). | 44.57 | 4.97 | 44.66 | 4.15 | 1.00 | 0.10 |
| AL | 23 – 24 | Cranial to caudal length of the acetabulum (7). | 20.21 | 1.05 | 21.48 | 1.15 | 1.00 | 0.06 |
| AW | 25 – 26 | Medial to lateral width of the acetabulum. | 19.02 | 1.26 | 20.12 | 1.2 | 1.00 | 0.07 |

**Table S3.** List and definitions of landmarks collected. Citations for the landmark definitions are indicated by a number referenced in the References at the end of this appendix.

| *No.* | *Landmark* | *Definition* |
|---|---|---|
| | | ***Basicranium*** |
| 1. | Inion | Location of the intersection of the right and left superior nuchal lines within the external occipital protuberance (8). |
| 2. | Opisthion | Midline point of the dorsal margin of the foramen magnum (8). |
| 3. | Basion | Midline point of the ventral margin of the foramen magnum (8). |
| 4. | Right margin of foramen magnum | Right margin of the foramen magnum at its greatest curvature (2). |
| 5. | Left margin of foramen magnum | Left margin of the foramen magnum at its greatest curvature (2). |
| 6. | Asterion | Point where the lambdoidal, parietomastoid, and occipitomastoid sutures meet (8). |
| 7. | Left mastoid | Inferior-most point of the mastoid process (8). |
| | | ***Scapula*** |
| 8. | Scapular spine lateral border | Point of intersection between the scapular spine and acromion process (3). |
| 9. | Scapular spine medial border | Medial margin of trapezius attachment along scapular spine (5). |
| 10. | Superior angle | Intersection of superior and medial borders of scapula (8). |
| 11. | Scapular spine vertebral border | Point on vertebral margin where the long axis of the scapular spine and vertebral border meet (5). |
| 12. | Inferior angle | Intersection of the lateral and medal borders of scapula (8). |
| 13. | Ventral margin of glenoid fossa | Ventral margin of glenoid fossa at its greatest width (5). |
| 14. | Dorsal margin of glenoid fossa | Dorsal margin of glenoid fossa at its greatest width (5). |
| 15. | Cranial margin of glenoid fossa | Superior-most margin of glenoid fossa (5). |
| 16. | Caudal margin of glenoid fossa | Inferior-most margin of glenoid fossa (5). |
| | | ***Clavicle*** |
| 17. | Acromial end | Lateral-most point of clavicle on cranial aspect. |
| 18. | Sternal end | Medial-most point of clavicle on cranial aspect. |
| | | ***Os coxa*** |
| 19. | Anterior superior iliac spine | Ventral-most point on lateral aspect of iliac crest (7). |
| 20. | Anterior inferior iliac spine | Ventral-most point on anterior inferior iliac spine. If only a bony roughening, the point is taken at the center of this rugosity (7). |
| 21. | Posterior superior iliac spine | Medial-most point of the dorsal-cranial border of iliac crest (7). |
| 22. | Posterior inferior iliac spine | Sharp projection posterior and inferior to the auricular surface (8). |
| 23. | Cranial margin of acetabulum | Cranial-most point of the margin of the acetabulum (7). |
| 24. | Caudal margin of acetabulum | Caudal-most point of the margin of the acetabulum, directly across from landmark 23 (7). |
| 25. | Medial margin of acetabulum | Medial-most point along margin of the acetabulum. |
| 26. | Lateral margin of acetabulum | Lateral-most point along margin of the acetabulum. |
| 27. | Cranial margin of auricular surface | Central point of the cranial-most margin of the auricular surface. |
| 28. | Caudal margin of auricular surface | Central point of the caudal-most margin of the auricular surface. |
| 29. | Center of acetabulum | Center of the acetabulum, defined as the intersection of the line between landmarks 23 and 24 and landmarks 27 and 28 (7). |

**Table S4.** Correlation matrix for all 21 traits. Highly correlated traits (r ≥ 0.8) are highlighted in orange.

| | **AW** | **AL** | **LIH** | **LIB** | **RAH** | **ASL** | **PMIB** | **AMIB** | **CLML** | **GL** | |
|---|---|---|---|---|---|---|---|---|---|---|---|
| **NR** | | 0.745 | 0.127 | 0.364 | 0.194 | 0.257 | 0.115 | 0.242 | 0.411 | 0.424 | **AW** |
| **SCM** | 0.187 | | 0.085 | 0.409 | 0.202 | 0.093 | 0.180 | 0.158 | 0.455 | 0.507 | **AL** |
| **SNL** | -0.012 | 0.242 | | -0.021 | -0.242 | -0.105 | 0.160 | -0.027 | 0.424 | 0.298 | **LIH** |
| **FMW** | -0.255 | 0.185 | 0.111 | | 0.440 | 0.231 | 0.207 | 0.301 | 0.582 | 0.437 | **LIB** |
| **FML** | -0.256 | 0.249 | 0.232 | 0.291 | | 0.250 | 0.565 | -0.018 | 0.298 | 0.132 | **RAH** |
| **SSL** | -0.029 | 0.163 | 0.057 | 0.374 | 0.162 | | -0.023 | -0.070 | 0.356 | 0.078 | **ASL** |
| **SBL** | 0.130 | 0.384 | 0.094 | 0.350 | 0.058 | 0.758 | | -0.017 | 0.498 | 0.269 | **PMIB** |
| **SBW** | -0.032 | 0.332 | 0.041 | 0.337 | 0.156 | 0.856 | 0.901 | | -0.041 | 0.101 | **AMIB** |
| **SBH** | 0.063 | 0.336 | 0.246 | 0.347 | 0.074 | 0.644 | 0.864 | 0.760 | | 0.579 | **CLML** |
| **VB** | 0.028 | 0.359 | 0.272 | 0.256 | 0.046 | 0.542 | 0.756 | 0.699 | 0.951 | | **GL** |
| **GW** | 0.058 | 0.405 | 0.123 | 0.266 | -0.032 | 0.644 | 0.572 | 0.628 | 0.562 | 0.507 | |
| **GL** | -0.022 | 0.374 | 0.233 | 0.411 | 0.192 | 0.535 | 0.439 | 0.483 | 0.584 | 0.523 | 0.602 |
| **CLML** | 0.207 | 0.473 | 0.047 | 0.456 | 0.118 | 0.684 | 0.774 | 0.705 | 0.702 | 0.584 | 0.699 |
| **AMIB** | -0.096 | -0.008 | 0.330 | -0.012 | 0.164 | 0.069 | 0.104 | 0.118 | 0.204 | 0.226 | -0.012 |
| **PMIB** | -0.090 | 0.055 | -0.072 | 0.005 | 0.192 | 0.302 | 0.286 | 0.333 | 0.269 | 0.268 | 0.305 |
| **ASL** | 0.189 | 0.566 | 0.144 | 0.222 | 0.061 | 0.051 | 0.189 | 0.160 | 0.123 | 0.126 | 0.304 |
| **RAH** | 0.085 | 0.177 | 0.202 | -0.046 | -0.069 | 0.004 | 0.073 | 0.051 | 0.179 | 0.289 | 0.219 |
| **LIB** | 0.358 | 0.292 | 0.165 | 0.321 | 0.053 | 0.493 | 0.536 | 0.440 | 0.614 | 0.597 | 0.362 |
| **LIH** | -0.080 | 0.159 | -0.147 | 0.152 | -0.099 | 0.338 | 0.371 | 0.271 | 0.219 | 0.068 | 0.383 |
| **AL** | 0.253 | 0.294 | 0.170 | 0.266 | -0.018 | 0.283 | 0.398 | 0.332 | 0.548 | 0.560 | 0.554 |
| **AW** | 0.241 | 0.277 | 0.208 | 0.255 | -0.078 | 0.171 | 0.246 | 0.164 | 0.355 | 0.371 | 0.334 |
| | **NR** | **SCM** | **SNL** | **FMW** | **FML** | **SSL** | **SBL** | **SBW** | **SBH** | **VB** | **GW** |

**Table S5. List of additional trait configurations.**

| *Configuration Designation* | *Basicranium* | *Shoulder Girdle* | *Pelvic Girdle* |
|---|---|---|---|
| 1 | NR, SCM, SNL, FMW, FML | SSL, SBL, VB, GW, CLML | AMIB, PMIB, ASL, LIH, AL |
| 2 | NR, SCM, SNL, FMW, FML | SSL, SBL, SBH, GW, CLML | AMIB, RAH, ASL, LIH, AL |
| 3 | NR, SCM, SNL, FMW, FML | SSL, SBL, VB, GL, CLML | AMIB, PMIB, ASL, LIH, AL |
| 4 | NR, SCM, SNL, FMW, FML | SSL, GL, VB, GW, CLML | AMIB, PMIB, ASL, LIH, AL |
| 5 | NR, SCM, SNL, FMW, FML | SSL, SBL, VB, GW, CLML | AMIB, PMIB, ASL, LIB, AL |
| 6 | NR, SCM, SNL, FMW, FML | SSL, SBL, VB, GW, CLML | AMIB, RAH, ASL, LIH, AL |
| 7 | NR, SCM, SNL, FMW, FML | SSL, SBL, VB, GW, CLML | AMIB, RAH, ASL, LIB, AL |
| 8 | NR, SCM, SNL, FMW, FML | SSL, SBL, GL, GW, CLML | AMIB, RAH, ASL, LIB, AL |
| 9 | NR, SCM, SNL, FMW, FML | SSL, SBL, VB, GL, CLML | AMIB, RAH, ASL, LIH, AL |
| 10 | NR, SCM, SNL, FMW, FML | SSL, SBL, VB, GW, CLML | AMIB, PMIB, ASL, LIH, AW |
| 11 | NR, SCM, SNL, FMW, FML | SSL, SBL, VB, GW, CLML | AMIB, RAH, ASL, LIB, AW |
| 12 | NR, SCM, SNL, FMW, FML | SSL, VB, GL, GW, CLML | AMIB, RAH, ASL, LIB, AL |

**Table S6.** Mean evolvability and 95% confidence intervals for each configuration of traits. Estimates for both sex-scaled and sex and size-scaled data is presented. Confidence intervals denoted with '~' indicate analyses that were singular. Traits comprising each configuration are listed in Table S3.

| *Configuration* | *Trait Groupings* | *Sex-scaled* | *Sex-scaled/Size-corrected* |
|---|---|---|---|
| 1 | All Traits | 0.0295 (0.0067, 0.0157) | 0.0241 (0.0035, 0.0095) |
| | Basicranium – Shoulder Girdle | 0.0285 (0.0074, 0.0175) | 0.0314 (0.0036, 0.0109) |
| | Shoulder Girdle – Pelvic Girdle | 0.0215 (0.0055, 0.0137) | 0.0213 (0.0025, 0.0081) |
| | Basicranium – Pelvic Girdle | 0.0302 (0.0065, 0.0163) | 0.0296 (0.0037, 0.0107) |
| 2 | All Traits | 0.0264 (0.0068, 0.0169) | 0.0072 (~) |
| | Basicranium – Shoulder Girdle | 0.0107 (0.0074, 0.0181) | 0.0084 (~) |
| | Shoulder Girdle – Pelvic Girdle | 0.0068 (0.0054, 0.0151) | 0.0047 (~) |
| | Basicranium – Pelvic Girdle | 0.0107 (0.0067, 0.0174) | 0.0089 (~) |
| 3 | All Traits | 0.0111 (0.0069, 0.0171) | 0.0063 (~) |
| | Basicranium – Shoulder Girdle | 0.0121 (0.0075, 0.0179) | 0.0068 (~) |
| | Shoulder Girdle – Pelvic Girdle | 0.0091 (0.0054, 0.0152) | 0.0153 (~) |
| | Basicranium – Pelvic Girdle | 0.0119 (0.0069, 0.0178) | 0.0079 (~) |
| 4 | All Traits | 0.0109 (0.0069, 0.0188) | 0.007 (0.0039, 0.0098) |
| | Basicranium – Shoulder Girdle | 0.0122 (0.0074, 0.0196) | 0.076 (0.0036, 0.0107) |
| | Shoulder Girdle – Pelvic Girdle | 0.0089 (0.0576, 0.0175) | 0.0048 (0.0029, 0.0081) |
| | Basicranium – Pelvic Girdle | 0.0118 (0.007, 0.0194) | 0.0089 (0.0041, 0.0111) |
| 5 | All Traits | 0.0107 (0.0069, 0.0164) | 0.0071 (~) |
| | Basicranium – Shoulder Girdle | 0.0127 (0.0075, 0.0179) | 0.0081 (~) |
| | Shoulder Girdle – Pelvic Girdle | 0.0089 (0.0051, 0.0145) | 0.0059 (~) |
| | Basicranium – Pelvic Girdle | 0.0113 (0.0069, 0.0171) | 0.0075 (~) |
| 6 | All Traits | 0.0127 (0.0071, 0.0183) | 0.0089 (0.0043, 0.0099) |
| | Basicranium – Shoulder Girdle | 0.0141 (0.0077, 0.0197) | 0.0098 (0.0042, 0.0112) |
| | Shoulder Girdle – Pelvic Girdle | 0.0108 (0.0055, 0.0167) | 0.0069 (0.0031, 0.0082) |
| | Basicranium – Pelvic Girdle | 0.0135 (0.0071, 0.0189) | 0.0104 (0.0042, 0.0112) |
| 7 | All Traits | 0.0178 (0.0068, 0.0174) | 0.0072 (~) |
| | Basicranium – Shoulder Girdle | 0.0192 (0.0077, 0.0186) | 0.0085 (~) |
| | Shoulder Girdle – Pelvic Girdle | 0.0164 (0.0051, 0.0154) | 0.0063 (~) |
| | Basicranium – Pelvic Girdle | 0.0187 (0.0068, 0.0181) | 0.0072 (~) |
| 8 | All Traits | 0.0103 (0.0068, 0.0156) | 0.0076 (~) |
| | Basicranium – Shoulder Girdle | 0.0122 (0.0076, 0.0171) | 0.0083 (~) |
| | Shoulder Girdle – Pelvic Girdle | 0.0088 (0.0049, 0.0137) | 0.0067 (~) |
| | Basicranium – Pelvic Girdle | 0.0105 (0.0068, 0.0163) | 0.0082 (~) |
| 9 | All Traits | 0.0126 (0.0072, 0.0189) | 0.0065 (0.0038, 0.0096) |
| | Basicranium – Shoulder Girdle | 0.0139 (0.0079, 0.0199) | 0.0078 (0.0037, 0.0109) |
| | Shoulder Girdle – Pelvic Girdle | 0.0108 (0.0056, 0.0172) | 0.0042 (0.0028, 0.0081) |
| | Basicranium – Pelvic Girdle | 0.0137 (0.0073, 0.0193) | 0.0085 (0.0037, 0.0106) |
| 10 | All Traits | 0.0102 (0.0069, 0.0172) | 0.0094 (~) |
| | Basicranium – Shoulder Girdle | 0.0117 (0.0074, 0.0185) | 0.0103 (~) |
| | Shoulder Girdle – Pelvic Girdle | 0.0081 (0.0056, 0.0154) | 0.0074 (~) |
| | Basicranium – Pelvic Girdle | 0.0108 (0.0069, 0.0176) | 0.0104 (~) |
| 11 | All Traits | 0.0091 (0.0066, 0.0155) | 0.0073 (~) |
| | Basicranium – Shoulder Girdle | 0.0112 (0.0074, 0.0169) | 0.0083 (~) |
| | Shoulder Girdle – Pelvic Girdle | 0.007 (0.0049, 0.0129) | 0.0064 (~) |
| | Basicranium – Pelvic Girdle | 0.0096 (0.0068, 0.0165) | 0.0077 (~) |
| 12 | All Traits | 0.0123 (0.0068, 0.0171) | 0.0078 (0.0046, 0.0098) |
| | Basicranium – Shoulder Girdle | 0.0137 (0.0077, 0.0184) | 0.0086 (0.0046, 0.0111) |
| | Shoulder Girdle – Pelvic Girdle | 0.0103 (0.0053, 0.0147) | 0.0064 (0.0033, 0.008) |
| | Basicranium – Pelvic Girdle | 0.0129 (0.0072, 0.0182) | 0.0088 (0.0049, 0.0113) |

**Table S7.** Conditioned covariance and 95% confidence intervals for each configuration of traits. Anatomical regions that are held constant by stabilizing selection are denoted by (x), whereas the anatomical regions under directional selection are denoted by (y).

| *Configuration* | *Trait Groupings* | *Sex-scaled* | *Sex-scaled/Size-corrected* |
|---|---|---|---|
| 1 | Basicranium (x) – Shoulder Girdle (y) | 0.0454 (0.016, 0.0667) | 0.0186 (0.0013, 0.0178) |
| | Shoulder Girdle (x) – Basicranium (y) | 0.068 (0.0222, 0.0881) | 0.0379 (0.0015, 0.0327) |
| | Basicranium (x) – Pelvic Girdle (y) | 0.0396 (0.0156, 0.0609) | 0.0248 (0.0014, 0.0234) |
| | Pelvic Girdle (x) – Basicranium (y) | 0.0724 (0.0268, 0.0934) | 0.0458 (0.0021, 0.044) |
| | Shoulder Girdle (x) – Pelvic Girdle (y) | 0.035 (0.0138, 0.0613) | 0.0229 (0.0013, 0.0217) |
| | Pelvic Girdle (x) – Shoulder Girdle (y) | 0.0411 (0.0178, 0.069) | 0.0160 (0.0015, 0.0225) |
| 2 | Basicranium (x) – Shoulder Girdle (y) | 0.0313 (0.017, 0.0734) | 0.0164 (0.0013, 0.0186) |
| | Shoulder Girdle (x) – Basicranium (y) | 0.0609 (0.0223, 0.0963) | 0.0344 (0.0016, 0.0367) |
| | Basicranium (x) – Pelvic Girdle (y) | 0.0330 (0149, 071) | 0.0230 (0.0015, 0.0232) |
| | Pelvic Girdle (x) – Basicranium (y) | 0.0672 (0.0277, 0.1001) | 0.0515 (0.0023, 0.0474) |
| | Shoulder Girdle (x) – Pelvic Girdle (y) | 0.0292 (0.0136, 0.0707) | 0.0165 (0.0013, 0.0212) |
| | Pelvic Girdle (x) – Shoulder Girdle (y) | 0.0286 (0.0173, 0.0756) | 0.0123 (0.0016, 0.0211) |
| 3 | Basicranium (x) – Shoulder Girdle (y) | 0.0394 (0.0152, 0.0747) | 0.0112 (0.001, 0.0163) |
| | Shoulder Girdle (x) – Basicranium (y) | 0.0632 (0.0219, 0.0949) | 0.0285 (0.0014, 0.0304) |
| | Basicranium (x) – Pelvic Girdle (y) | 0.0409 (0.016, 0.0723) | 0.0163 (0.0012, 0.0217) |
| | Pelvic Girdle (x) – Basicranium (y) | 0.0692 (0.0139, 0.0975) | 0.0408 (0.0019, 0.0409) |
| | Shoulder Girdle (x) – Pelvic Girdle (y) | 0.0352 (0.0139, 0.072) | 0.0116 (0.001, 0.0216) |
| | Pelvic Girdle (x) – Shoulder Girdle (y) | 0.0376 (0.0156, 0.0761) | 0.0075 (0.0013, 0.0209) |
| 4 | Basicranium (x) – Shoulder Girdle (y) | 0.0383 (0.0181, 0.0845) | 0.0141 (0.0018, 0.0208) |
| | Shoulder Girdle (x) – Basicranium (y) | 0.0634 (0.0246, 0.1036) | 0.0330 (0.0022, 0.039) |
| | Basicranium (x) – Pelvic Girdle (y) | 0.0371 (0.0172, 0.0821) | 0.0252 (0.0021, 0.0254) |
| | Pelvic Girdle (x) – Basicranium (y) | 0.0668 (0.0284, 0.1049) | 0.0382 (0.003, 0.0466) |
| | Shoulder Girdle (x) – Pelvic Girdle (y) | 0.0314 (0.0156, 0.0834) | 0.0191 (0.0018, 0.0246) |
| | Pelvic Girdle (x) – Shoulder Girdle (y) | 0.0332 (0.0187, 0.0857) | 0.0118 (0.0018, 0.0218) |

Table S7 Continued

| | | | |
|---|---|---|---|
| 5 | Basicranium (x) – Shoulder Girdle (y) | 0.0404 (0.015, 0.0717) | 0.0162 (0.0015, 0.0194) |
| | Shoulder Girdle (x) – Basicranium (y) | 0.0609 (0.0245, 0.0965) | 0.0194 (0.0023, 0.0385) |
| | Basicranium (x) – Pelvic Girdle (y) | 0.0334 (0.0142, 0.0661) | 0.0072 (0.0019, 0.0231) |
| | Pelvic Girdle (x) – Basicranium (y) | 0.0651 (0.0271, 0.0969) | 0.0241 (0.0028, 0.0449) |
| | Shoulder Girdle (x) – Pelvic Girdle (y) | 0.0294 (0.0123, 0.0664) | 0.0138 (0.0018, 0.0222) |
| | Pelvic Girdle (x) – Shoulder Girdle (y) | 0.0392 (0.0147, 0.0719) | 0.0118 (0.0116, 0.0222) |
| 6 | Basicranium (x) – Shoulder Girdle (y) | 0.0519 (0.0185, 0.0819) | 0.0177 (0.0016, 0.0202) |
| | Shoulder Girdle (x) – Basicranium (y) | 0.0756 (0.024, 0.1029) | 0.0368 (0.0018, 0.0388) |
| | Basicranium (x) – Pelvic Girdle (y) | 0.0473 (0.0171, 0.0786) | 0.0253 (0.0018, 0.0252) |
| | Pelvic Girdle (x) – Basicranium (y) | 0.0782 (0.0306, 0.1037) | 0.0534 (0.0026, 0.0486) |
| | Shoulder Girdle (x) – Pelvic Girdle (y) | 0.0446 (0.0147, 0.0789) | 0.0238 (0.0014, 0.0238) |
| | Pelvic Girdle (x) – Shoulder Girdle (y) | 0.0492 (0.0188, 0.083) | 0.0120 (0.0017, 0.0222) |
| 7 | Basicranium (x) – Shoulder Girdle (y) | 0.0819 (0.0164, 0.0754) | 0.0163 (0.0012, 0.0191) |
| | Shoulder Girdle (x) – Basicranium (y) | 0.1030 (0.023, 0.0999) | 0.0205 (0.0018, 0.0374) |
| | Basicranium (x) – Pelvic Girdle (y) | 0.0777 (0.0139, 0.0708) | 0.0084 (0.0014, 0.0225) |
| | Pelvic Girdle (x) – Basicranium (y) | 0.1031 (0.0277, 0.1004) | 0.0181 (0.0019, 0.0415) |
| | Shoulder Girdle (x) – Pelvic Girdle (y) | 0.0774 (0.0116, 0.0715) | 0.0166 (0.0014, 0.0205) |
| | Pelvic Girdle (x) – Shoulder Girdle (y) | 0.0816 (0.0166, 0.0775) | 0.0125 (0.0012, 0.021) |
| 8 | Basicranium (x) – Shoulder Girdle (y) | 0.0415 (0.0156, 0.0668) | 0.0201 (0.0016, 0.0191) |
| | Shoulder Girdle (x) – Basicranium (y) | 0.0554 (0.0221, 0.0909) | 0.0318 (0.0033, 0.0375) |
| | Basicranium (x) – Pelvic Girdle (y) | 0.0267 (0.0144, 0.0626) | 0.0121 (0.0018, 0.0236) |
| | Pelvic Girdle (x) – Basicranium (y) | 0.0584 (0.0259, 0.0916) | 0.0200 (0.0032, 0.0462) |
| | Shoulder Girdle (x) – Pelvic Girdle (y) | 0.0231 (0.0124, 0.0625) | 0.0157 (0.0018, 0.0218) |
| | Pelvic Girdle (x) – Shoulder Girdle (y) | 0.0369 (0.0153, 0.0671) | 0.0193 (0.0016, 0.021) |

Table S7 Continued

| | | | |
|---|---|---|---|
| 9 | Basicranium (x) – Shoulder Girdle (y) | 0.0495 (0.0185, 0.0842) | 0.0132 (0.0012, 0.0171) |
| | Shoulder Girdle (x) – Basicranium (y) | 0.0740 (0.0248, 0.1072) | 0.0302 (0.0016, 0.0345) |
| | Basicranium (x) – Pelvic Girdle (y) | 0.0497 (0.0179, 0.0804) | 0.0153 (0.0013, 0.0235) |
| | Pelvic Girdle (x) – Basicranium (y) | 0.0773 (0.0317, 0.1106) | 0.0388 (0.002, 0.042) |
| | Shoulder Girdle (x) – Pelvic Girdle (y) | 0.0471 (0.0168, 0.0803) | 0.0135 (0.0011, 0.0215) |
| | Pelvic Girdle (x) – Shoulder Girdle (y) | 0.0485 (0.0195, 0.0856) | 0.0077 (0.0012, 0.022) |
| 10 | Basicranium (x) – Shoulder Girdle (y) | 0.0333 (0.0176, 0.0762) | 0.0186 (0.0014, 0.0179) |
| | Shoulder Girdle (x) – Basicranium (y) | 0.0577 (0.0223, 0.0961) | 0.0384 (0.0019, 0.0359) |
| | Basicranium (x) – Pelvic Girdle (y) | 0.0308 (0.0171, 0.0716) | 0.0238 (0.0016, 0.0245) |
| | Pelvic Girdle (x) – Basicranium (y) | 0.0629 (0.0284, 0.0993) | 0.0482 (0.002, 0.0483) |
| | Shoulder Girdle (x) – Pelvic Girdle (y) | 0.0229 (0.0146, 0.0702) | 0.0239 (0.0015, 0.0221) |
| | Pelvic Girdle (x) – Shoulder Girdle (y) | 0.0264 (0.0181, 0.0779) | 0.0178 (0.0015, 0.021) |
| 11 | Basicranium (x) – Shoulder Girdle (y) | 0.0289 (0.0143, 0.0623) | 0.0178 (0.0011, 0.018) |
| | Shoulder Girdle (x) – Basicranium (y) | 0.0467 (0.021, 0.0904) | 0.0314 (0.0019, 0.0387) |
| | Basicranium (x) – Pelvic Girdle (y) | 0.0185 (0.013, 0.0573) | 0.0122 (0.0014, 0.0226) |
| | Pelvic Girdle (x) – Basicranium (y) | 0.0489 (0.0254, 0.088) | 0.0184 (0.0019, 0.0427) |
| | Shoulder Girdle (x) – Pelvic Girdle (y) | 0.0109 (0.0108, 0.0574) | 0.0148 (0.0014, 0.0217) |
| | Pelvic Girdle (x) – Shoulder Girdle (y) | 0.0200 (0.014, 0.0635) | 0.0108 (0.0012, 0.0207) |
| 12 | Basicranium (x) – Shoulder Girdle (y) | 0.0492 (0.015, 0.0702) | 0.0211 (0.0023, 0.0214) |
| | Shoulder Girdle (x) – Basicranium (y) | 0.0721 (0.0231, 0.0992) | 0.0318 (0.0038, 0.0436) |
| | Basicranium (x) – Pelvic Girdle (y) | 0.0436 (0.0134, 0.0673) | 0.0109 (0.0028, 0.0242) |
| | Pelvic Girdle (x) – Basicranium (y) | 0.0700 (0.0258, 0.0985) | 0.0173 (0.0033, 0.0413) |
| | Shoulder Girdle (x) – Pelvic Girdle (y) | 0.0415 (0.012, 0.0676) | 0.0183 (0.0024, 0.024) |
| | Pelvic Girdle (x) – Shoulder Girdle (y) | 0.0456 (0.0159, 0.0718) | 0.0181 (0.0022, 0.0214) |

**Table S8.** Mean evolvability and 95% confidence intervals for macaques (Macaca mulatta group) and tamarins (*Saguinus oedipus* and *Saguinus fuscicollis illigeri*). Estimates for both sex-scaled and sex and size-scaled data are presented.

| ***Taxon*** | ***Trait Groupings*** | ***Sex-scaled*** | ***Sex-scaled/Size-corrected*** |
|---|---|---|---|
| *Macaca* | *All Traits* | 0.0121 (0.0073, 0.0179) | 0.0097 (0.0049, 0.0111) |
| | *Basicranium – Shoulder Girdle* | 0.0123 (0.0076, 0.0184) | 0.0097 (0.005, 0.0119) |
| | *Shoulder Girdle – Pelvic Girdle* | 0.0129 (0.007, 0.0185) | 0.0098 (0.0046, 0.0119) |
| | *Basicranium – Pelvic Girdle* | 0.0115 (0.007, 0.0176) | 0.0095 (0.0043, 0.0112) |
| *Saguinus* | *All Traits* | 0.0046 (0.0029, 0.0059) | 0.0029 (0.0018, 0.0038) |
| | *Basicranium – Shoulder Girdle* | 0.0054 (0.0031, 0.0065) | 0.0036 (0.0018, 0.0044) |
| | *Shoulder Girdle – Pelvic Girdle* | 0.0044 (0.0027, 0.0058) | 0.0026 (0.0016, 0.0037) |
| | *Basicranium – Pelvic Girdle* | 0.0041 (0.0026, 0.0058) | 0.0027 (0.0016, 0.0038) |

**Table S9.** Mean conditioned covariance and 95% confidence intervals for macaques (Macaca mulatta group, n=72) and tamarins (*Saguinus oedipus* and *Saguinus fuscicollis illigeri*, n=43). Estimates for both sex-scaled and sex and size-scaled data are presented.

| ***Taxon*** | ***Trait Groupings*** | ***Sex-scaled*** | ***Sex-scaled/Size-corrected*** |
|---|---|---|---|
| *Macaca* | *Basicranium (x) – Shoulder Girdle (y)* | 0.0483 (0.0197, 0.087) | 0.0257 (0.0029, 0.0304) |
| | *Shoulder Girdle (x) – Basicranium (y)* | 0.0426 (0.0182, 0.079) | 0.0245 (0.0022, 0.028) |
| | *Basicranium (x) – Pelvic Girdle (y)* | 0.0487 (0.0201, 0.0816) | 0.0230 (0.0029, 0.0259) |
| | *Pelvic Girdle (x) – Basicranium (y)* | 0.0480 (0.022, 0.0828) | 0.0281 (0.0028, 0.0317) |
| | *Shoulder Girdle (x) – Pelvic Girdle (y)* | 0.0397 (0.0152, 0.074) | 0.0128 (0.0019, 0.0198) |
| | *Pelvic Girdle (x) – Shoulder Girdle (y)* | 0.0475 (0.0166, 0.0832) | 0.0150 (0.0025, 0.0289) |
| *Saguinus* | *Basicranium (x) – Shoulder Girdle (y)* | 0.0249 (0.0085, 0.0293) | 0.0127 (0.0023, 0.0134) |
| | *Shoulder Girdle (x) – Basicranium (y)* | 0.0216 (0.0073, 0.029) | 0.0146 (0.0018, 0.014) |
| | *Basicranium (x) – Pelvic Girdle (y)* | 0.0145 (0.0069, 0.0238) | 0.0054 (0.002, 0.01) |
| | *Pelvic Girdle (x) – Basicranium (y)* | 0.0231 (0.0099, 0.0306) | 0.0123 (0.0027, 0.0159) |
| | *Shoulder Girdle (x) – Pelvic Girdle (y)* | 0.0139 (0.006, 0.0232) | 0.0043 (0.0017, 0.0092) |
| | *Pelvic Girdle (x) – Shoulder Girdle (y)* | 0.0250 (0.0092, 0.0299) | 0.0095 (0.0023, 0.0143) |

**Table S10.** Mean evolvability for pairwise comparisons of anatomical groupings using 20 traits evenly distributed across the following four anatomical regions: basicranium, shoulder girdle, pelvic girdle, and humerus. Traits for the basicranium, shoulder girdle, and pelvic girdle are described in the main text. The five traits from the humerus are measures of maximum length, width of the distal joint surface, distance between the medial and lateral epi-condyles, distance from the medial epicondyle to medial border of distal joint surface, and the distance from the lateral epicondyle to the lateral border of the distal joint surface. Abbreviations for pairings of anatomical regions are as follow: basicranium and pelvic girdle (B-PG), basicranium and shoulder girdle (B-SG), shoulder girdle and pelvic girdle (SG-PG), basicranium and humerus (B-H), pelvic girdle and humerus (PG-H), shoulder girdle and humerus (SG-H).

| *Anatomical Regions* | **B - PG** | **B - SG** | **SG - PG** | **B - H** | **PG - H** | **SG - H** |
|---|---|---|---|---|---|---|
| *Mean* | 0.0116 | 0.0094 | 0.0097 | 0.0129 | 0.0134 | 0.0109 |

**References**


1. Richmond, F.J.R., Singh, K., & Corneil, B.D. (2001). Neck Muscles in the Rhesus Monkey. I. Muscle Morphometry and Histochemistry. *Journal of Neurophysiology* **86**, 1717-1728.

2. Moore-Jansen, P.M., Ousley, S.D., & Jantz, R.J. (1994). *Data Collection Procedures for Forensic Skeletal Material*. Report of Investigations No. 48. Knoxville, TN: The University of Tennessee.

3. Sears, K.E., Bianchi, C., Powers, L., & Beck, A.L. (2013). Integration of the mammalian shoulder girdle within populations and over evolutionary time. *Journal of Evolutionary Biology* **26**, 1536-1548.

4. Martin, R. (1928). *Lehrbuch der Anthropologie in Systematischer Darstellung mit Besonderer Berücksichtigung der Anthropologischen Methoden für Studierende Ärtze und Forschungsreisende.* Zweiter Band: Kraniologie, Osteologie. Second Edition. Jena: Gustav Fischer.

5. Young, N.M. (2006). Function, ontogeny and canalization of shape variance in the primate scapula. *Journal of Anatomy* **209**, 623-636.

6. Grabowski, M., & Roseman, C.C. (2015). Complex and changing patterns of natural selection explain evolution of the human hip. *Journal of Human Evolution* **85**, 94-110.

7. Lewton, K.L. (2012). Evolvability of the primate pelvic girdle. *Evolutionary Biology* **39**, 126-139.

8. White, T.D., Black, M.T., &Folkens P.A. (2012). *Human Osteology.* Third Edition. New York: Academic Press.
